# Genomic features of *Mycobacterium avium* subsp. *hominissuis* isolated from pigs in Japan

**DOI:** 10.1101/2021.06.15.447579

**Authors:** Tetsuya Komatsu, Kenji Ohya, Atsushi Ota, Yukiko Nishiuchi, Hirokazu Yano, Kayoko Matsuo, Justice Opare Odoi, Shota Suganuma, Kotaro Sawai, Akemi Hasebe, Tetsuo Asai, Tokuma Yanai, Hideto Fukushi, Takayuki Wada, Shiomi Yoshida, Toshihiro Ito, Kentaro Arikawa, Mikihiko Kawai, Manabu Ato, Anthony D Baughn, Tomotada Iwamoto, Fumito Maruyama

## Abstract

*Mycobacterium avium* subsp. *hominissuis* (MAH) is one of the most important agents causing non-tuberculosis mycobacterial infection in humans and pigs. Genome analysis on MAH of human isolates has been proceeding, however, those of pigs are limited despite its potential source of infection to human. In the current study, we obtained 30 draft genome sequences of MAH of pigs reared in Japan. The 30 draft genomes consisted of 4,848,678 – 5,620,788 bp length, 4,652 – 5,388 coding genes and 46 – 75 (Med: 47) tRNAs. All isolates had restriction modification associated genes and 185 – 222 predicted virulence genes. Two isolates had tRNA arrays and one isolate had a clustered regularly interspaced short palindromic repeat (CRISPR) region. Our results will be useful for evaluation of the ecology of MAH by providing a foundation for genome-based epidemiological studies.

## Data description

### Context

To date, incidence of infection caused by non-tuberculous mycobacteria (NTM) has been increasing all over the world [1]. Among NTMs, *Mycobacterium avium* complex (MAC) is one of the most critical agents. MAC has 4 subspecies, namely *M. avium* subsp. *avium* (MAA), *M. avium* subsp. *paratuberculosis* (MAP), *M. avium* subsp. *silvaticum* (MAS) and *M. avium* subsp. *hominissuis* (MAH). MAH is known as a major pathogen for humans, causing lung disease and sometimes disseminated infection in immune suppressed patients [2, 3]. MAH is also a main causative agent of mycobacteriosis in pigs [4], showing mesenteric and mandibular lymphadenitis [5] and sometimes systemic infection [6]. Swine mycobacteriosis exerts severe economic impact in affected farms. MAH infected pigs are suspected as potential risk for human infection [7, 8, 9, 10].

Recently, genomic epidemiological study of MAH has extensively progressed. In our recent studies, MAH is divided into 6 major lineages (MahEastAsia1, MahEastAsia2, SC1 - SC4) and each lineage is predominant in specific regions on a global scale [11, 12]. For example, the MahEastAsia1 and MahEastAsia2 are frequently isolated from human lung disease in Japan and Korea although SC1 – 4 are isolated from America and Europe [11, 12]. Japanese pig isolates are mainly classified into 2 lineages, SC2 and SC4 [11, 12]. However, from the one health point of view, to exactly clarify the ecology of MAH, the number of pig isolates used in these studies was insufficient.

As stated above, genome-based analysis of MAH has been proceeding and the most essential genes of MAH are thought to be mutual orthologues of genes in *Mycobacterium tuberculosis* (MTB) [13]. Although components of virulence systems have been investigated [14], reports about genome contents, even drug resistance genes are not available, despite the increasing the incidence of MAH disease [1]. To understand MAH evolution, distribution and to promote the identification of targets for antimicrobial drug discovery, the characterization of the defining genomic features of MAH is essential.

Here we obtained draft genome sequences of 30 MAH isolates derived from pigs reared in Japan, and identified genome features for bacterial defense systems, such as restriction modification (RM) system, clustered regularly interspaced short palindromic repeat (CRISPR), tRNA arrays, virulence factors and drug resistance genes. Our results in this study may provide a way to understand the epidemiological relationship of MAH in human and pigs.

## Methods

### a) Sampling

MAH isolates were collected from pigs reared at two areas, Tokai and Hokuriku in Japan, where about 10 % of pigs in Japan are reared. 48 mesenteric or mandibular lymph nodes of pigs reared in Tokai area were collected from Gifu meat inspection center from July – December, 2015. Samples (20: mesenteric lymph nodes, 1: mandibular lymph nodes, 1: liver) of Tokai and Hokuriku area were collected between August, 1998 – Mar, 2018 and archived in Toyama meat inspection center.

### b) Bacterial isolation and DNA extraction

The method of bacterial isolation was available in protocols. io [15]. The mesenteric or mandibular lymph nodes with mycobacterial granulomatous lesions were mixed with 400ul of 2% NaOH and incubated at room temperature overnight. The samples were spread onto 2% Ogawa medium (Kyokuto Pharmaceutical, Tokyo, Japan) and incubated at 37 °C for 3 – 4 weeks. A single colony was inoculated onto 7H11 broth with 10% oleic acid-albumin-dextrose-catalase as a supplement. The isolates were stored with Microbank (Pro Lab Diagnostics Inc., Richmond Hill, ON, Canada) at -80°C. The method of extraction of genomic DNA was also available in protocols. io [16]. In brief, cells were delipidated by treatment with acetone, then lysed by lysozyme and Proteinase K. Genomic DNA was extracted by phenol/chloroform treatment of the lysates.

### c) Identification of MAH and insertion sequence profile

PCR amplification of *M. avium* 16S rRNA genes (MAV) was conducted for screening [17]. Isolates positive for MAV were identified by sequencing *hsp65* and *rpoB* genes [18, 19]. Basic Local Alignment Search Tool (BLAST) analysis was conducted using partial sequences of *rpoB* gene. Phylogenetic analysis of both genes was conducted by maximum likelihood method using Molecular Evolutionary Genetics Analysis (MEGA) software ver. 7.0. Bootstrap values were calculated from 1,000 replications. Insertion sequence patterns of IS*900*, IS*901*, IS*902* and IS*1245* were performed as described previously [20, 21, 22]. IS*1311* and IS*1613* were searched for within draft genomes by using ISfinder (https://isfinder.biotoul.fr) with default parameters [23].

### d) Draft genome sequences and genome annotation

Extraction of genomic DNA was described above. An average 350-bp paired-end libraries were prepared from extracted genomic DNA by TruSeq DNA PCR-Free High Throughput Library Prep Kit (Illumina, San Diego, CA, USA). Pair-end sequencing (2× 150-bp) was conducted using the HiSeq X Ten sequencing platform (Illumina) at the Beijing Genomics Institute (Shenzhen, China). Output reads were trimmed by TrimGalore! (https://github.com/FelixKrueger/TrimGalore) and were corrected its mismatched reads by SPAdes ver 3.12.0. [24]. The reads were assembled and polished using Pilon [25] and Unicycler [26], and then genome completeness was estimated by CheckM [27]. Taxonomic classification of contigs was carried out using Kaiju [28] and Anvi’o [29]. Draft genome sequences were annotated via the National Center for Biotechnology Information (NCBI) Prokaryotic Genome Annotation Pipeline (PGAP) [30].

### e) Detection of bacterial defence systems (RM system and CRISPR CAS system) in MAH genome

RM systems were determined by online tool, Restriction-ModificationFinder version 1.1 (https://cge.cbs.dtu.dk/services/Restriction-ModificationFinder/) twice with the following settings (1: database: All incl. putative genes, threshold for %ID: 90%, minimum length: 80% to search the RM system of MAH and 2: database: All, threshold for %ID: 10%, minimum length: 20% to confirm the orthologue of MTB or the other Mycobacteria) [31]. CRISPR Cas systems were identified by the online tool CRISPRCasFinder program (https://crisprcas.i2bc.paris-saclay.fr/CrisprCasFinder/Index) with default setting [32, 33].

### f) Detection of tRNA arrays in MAH genome

Total number of tRNAs in this study were retrieved from gb files annotated by PGAP. Draft genomes of GM17 and OCU479 isolates, which had more tRNAs than the others (Table 1), were inspected by tRNAscan-SE (http://lowelab.ucsc.edu/tRNAscan-SE/) to search tRNA arrays [34]. tRNA gene isotype synteny (expressed by the single-letter amino acid code) of both isolates and the reference strains were aligned and used for the maximum likelihood method by MEGA 7.0. Classification of both isolates was conducted as previously described [35].

**Table 1.** Summary information for the draft genome sequences of 30 MAH isolates in this study. * CDSs: coding sequences.

| Isolate | Genome size<br>(bp) | N50<br>(bp) | Coverage | No. of<br>contig | G+C<br>content (%) | No. of<br>CDSs* | No. of<br>tRNAs |
| --- | --- | --- | --- | --- | --- | --- | --- |
| GM5 | 5,037,010 | 35,760 | 277 | 224 | 69.06 | 4,877 | 47 |
| GM10 | 4,858,055 | 33,212 | 277 | 248 | 69.16 | 4,708 | 47 |
| GM12 | 4,848,678 | 33,219 | 253 | 261 | 69.17 | 4,732 | 47 |
| GM16 | 5,012,047 | 24,262 | 274 | 346 | 68.84 | 4,981 | 46 |
| GM17 | 5,265,075 | 30,906 | 355 | 289 | 68.77 | 5,190 | 75 |
| GM21 | 4,899,737 | 45,080 | 411 | 216 | 69.20 | 4,734 | 47 |
| GM32 | 4,897,271 | 47,147 | 292 | 208 | 69.20 | 4,712 | 47 |
| GM44 | 5,086,547 | 26,307 | 251 | 316 | 68.95 | 4,780 | 46 |
| OCU467 | 5,110,693 | 243,182 | 207 | 75 | 69.16 | 4,803 | 46 |
| OCU468 | 5,459,638 | 137,464 | 198 | 132 | 68.96 | 5,176 | 46 |
| OCU469 | 5,167,480 | 190,329 | 191 | 57 | 69.19 | 4,886 | 47 |
| OCU470 | 5,388,572 | 124,661 | 220 | 132 | 68.98 | 5,103 | 46 |
| OCU471 | 4,990,913 | 193,095 | 237 | 70 | 69.24 | 4,713 | 47 |
| OCU472 | 5,410,552 | 119,264 | 180 | 139 | 68.97 | 5,163 | 47 |
| OCU473 | 5,237,229 | 105,027 | 232 | 118 | 69.11 | 4,981 | 47 |
| OCU474 | 5,087,878 | 168,670 | 213 | 81 | 69.26 | 4,817 | 47 |
| OCU475 | 5,376,580 | 113,114 | 243 | 130 | 68.99 | 5,121 | 46 |
| OCU476 | 5,359,545 | 133,302 | 268 | 132 | 69.00 | 5,094 | 46 |
| OCU477 | 5,087,664 | 218,065 | 221 | 85 | 69.22 | 4,779 | 47 |
| OCU478 | 5,108,303 | 272,265 | 230 | 73 | 69.17 | 4,803 | 46 |
| OCU479 | 5,620,788 | 112,152 | 167 | 143 | 68.78 | 5,388 | 75 |
| OCU480 | 5,088,946 | 195,446 | 53 | 73 | 69.24 | 4,820 | 47 |
| OCU481 | 5,100,722 | 163,519 | 247 | 101 | 69.19 | 4,802 | 47 |
| OCU482 | 5,100,769 | 163,705 | 244 | 99 | 69.19 | 4,800 | 47 |
| OCU483 | 4,943,024 | 200,611 | 228 | 68 | 69.24 | 4,652 | 47 |
| OCU484 | 5,096,430 | 141,792 | 249 | 104 | 69.20 | 4,811 | 47 |
| OCU485 | 5,109,020 | 243,182 | 258 | 80 | 69.16 | 4,805 | 46 |
| OCU486 | 5,023,805 | 234,302 | 40 | 52 | 69.23 | 4,722 | 47 |
| Toy194 | 5,347,524 | 216,164 | 273 | 93 | 68.97 | 5,018 | 47 |
| Toy195 | 5,346,468 | 168,809 | 192 | 103 | 68.97 | 5,029 | 47 |

### g) Detection of virulence factors and drug resistance genes

Virulence genes were identified by using VFanalyzer (http://www.mgc.ac.cn/VFs/main.htm) [36]. We selected the following settings, genus: Mycobacterium, specify a representative genome: *M. avium* 104 and choose genomes for comparison: blank and draft genome fasta files were uploaded. Drug resistance genes were identified by Resistance Gene Identifier (RGI) version 5.1.0 (https://card.mcmaster.ca/analyze/rgi) with the following settings, Select Data Type: DNA sequence, Select Criteria: Perfect and Strict hit only, Nudge ≥95% identity Loose hits to Strict: Exclude nudge, Sequence Quality: high quality/coverage [37]. To confirm the existence of mutations detected by RGI, we retrieved the respective drug resistance associated genes from draft genome sequences, aligned by MEGA 7.0., and then manually checked for mutations in the nucleotide sequences.

## Data Validation and quality control

### Identification of MAH

The experimental workflow from sampling to identification is shown in Fig. 1. We successfully obtained 13 MAH isolates derived from Tokai area and 8 out of 13 isolates (GM5 – GM44) with 22 isolates of Tokai and Hokuriku area (OCU467 – OCU486, Toy194, Toy195) were used for draft genome sequence analysis. We conducted multiple examinations to determine the isolates as MAH, IS possession patterns, sequence analysis of *hsp65* (Supplementary Table 1). Among MAH subspecies, the patterns of IS possession is different and is used for subspecies identification [38]. IS*900* and IS*901* are known as the indicator of MAP and MAA, respectively [21, 22]. MAH is usually positive for IS*1245* [39], and is negative for IS*900*, IS*901* and IS*902* [20], however, MAH strains without IS*1245* are frequently distributed in Japan [39, 40]. In our study, 10/30 isolates were negative for IS*1245* (33.3%) and none had IS*900*, IS*901* and IS*902* (Supplementary Table 1). In general, subspecies of *M. avium* is also identified by *hsp65* gene analysis, which had 17 variations of SNP among subspecies [19]. MAH has usually 1, 2, 3, 7, 8 or 9 *hsp* code [19], however, five isolates had unclassified *hsp* code (indicated by N) in this study (Supplementary Table 1). Therefore, we also conducted partial sequence analysis of the *rpoB* gene and the isolates were identified as MAH by BLAST analysis. In addition, we conducted phylogenetic analysis based on *hsp65* and *rpoB* genes retrieved from draft genome and all isolates in this study were also classified into MAH (Fig 2). All of these examinations confirmed that our isolates were MAH.

**Figure 1.**
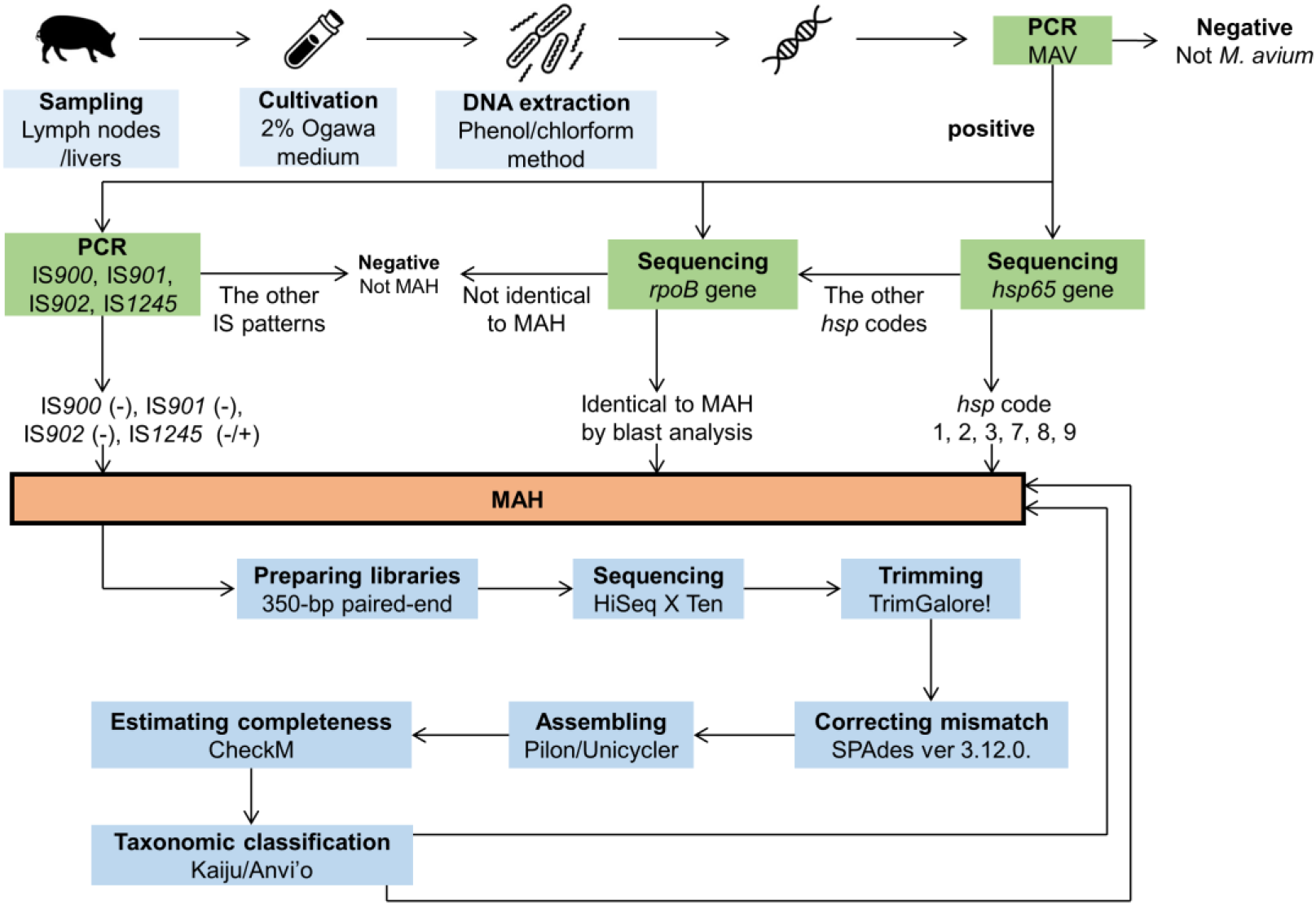
The experimental workflows in this study.

**Figure 2.**
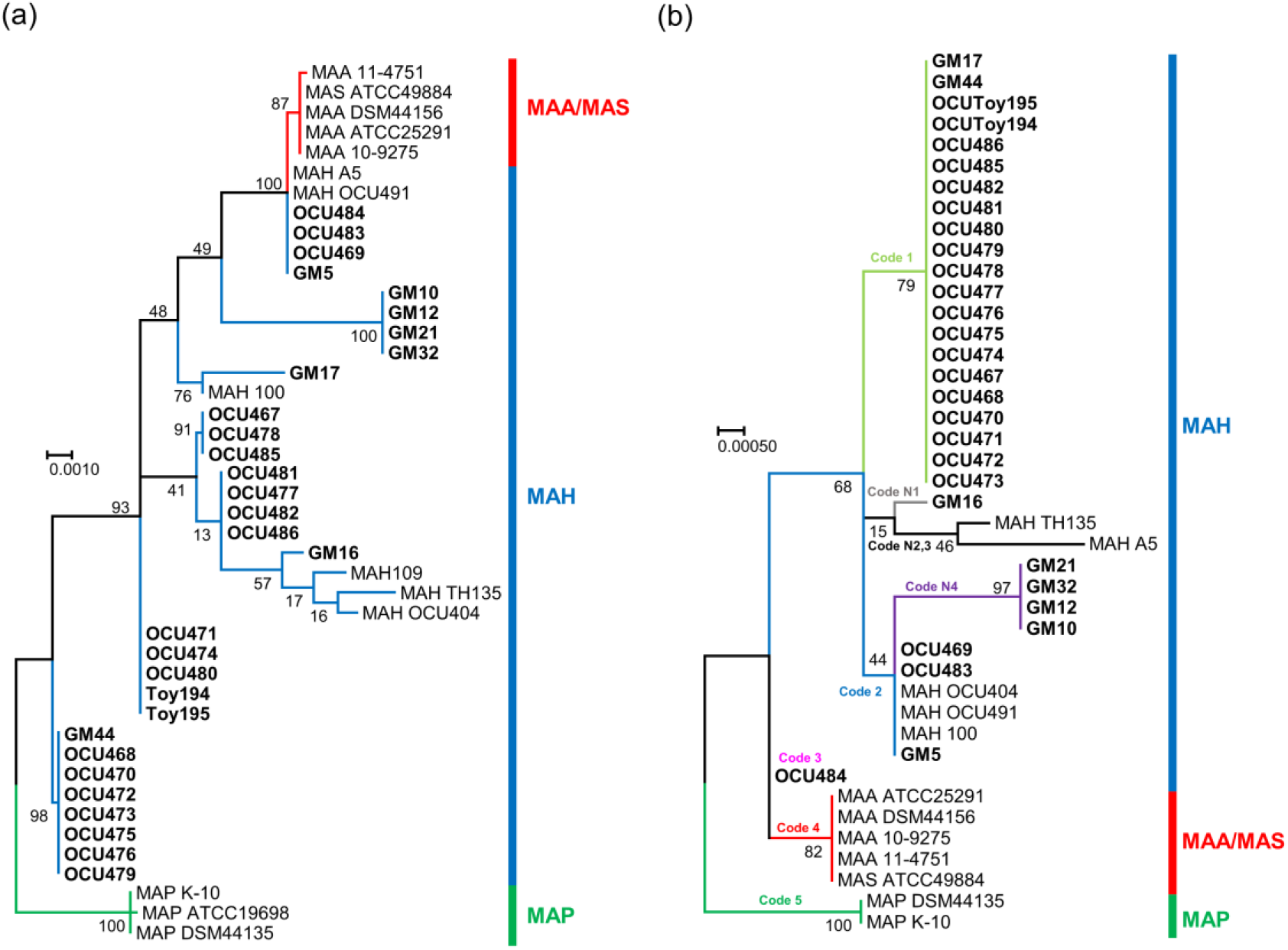
Phylogenetic analysis based on *rpoB* gene and *hsp65* gene. Phylogenetic tree was generated by maximum likelihood method using MEGA 7.0. All isolates in this study are indicated in bold font. **(a)** 30 MAH isolates in this study were classified as MAH and were differentiated from MAP and MAA/MAS node. **(b)** All the isolates in this study were classified into 5 *hsp* code, code 1, 2, 3, N1 and N4. These isolates were differentiated from MAP and MAA/MAS nodes. The bootstrap values were determined from 1,000 replications. The scale bar indicates genetic distances among strains.

### Draft genome data

All of our draft genome sequences had a total length between 4.85 – 5.62 Mb, similar to complete MAH genomes [41, 42]. All isolates had over 24kb N50 and over 40 fold genome coverage (average 233) (Table 1).

### Genome content analysis

In total, we identified 73 putative RM systems, including 24 type I RM systems, 48 type II RM systems, and 1 type III RM systems (Supplementary Table 2). All isolates had at least one Type II RM system and GM5, GM16, GM17, OCU468 – OCU470, OCU472, OCU473, OCU475, OCU476, OCU479, OCU483 and OCU484 had Type I, Type II RM systems, and GM44 had 3 types of RM systems. In these RM systems, 7 RM systems had homologues in MTB and 30 RM systems had homologues in *M. kansasii*. Orphan methyltransferase was detected in OCU473 and OCU479. CRISPR was detected only in GM44 (Supplementary Table 3). The sequences of the region were identical to MAH 104 (Query Cover: 100%, E value: 0.0, Per. Ident: 99.99%) which is the only MAH strain that had an intact CRISPR in the database (https://crisprcas.i2bc.paris-saclay.fr/MainDb/StrainList). The isolates had 185 – 222 virulence factors and 141 factors were common in all isolates (Supplementary Table 4). All isolates shared the same 2 drug resistance genes, *mtrA* which is associated with cell division and cell wall integrity [43] and resistance to macrolide antibiotics, and *RbpA* which regulates bacterial transcription and is associated with rifampicin resistance (Supplementary Table 5) [44]. In addition, single nucleotide polymorphisms (SNP) associated with drug resistance were found. All isolates had a C117D change in the *murA* gene conferring resistance to fosfomycin. A A2274G mutation in the *Mycobacterium avium* 23S rRNA which contributes to macrolide resistance was also detected by RGI, but when we examined the aligned nucleotide sequence, no point mutation was found in all isolates (Supplementary Table 5). CRISPR, virulence factor and drug resistance genes were selected from online tools. Original databases of each tool used in this study were updated in 2020, suggesting our data are based on the forefront of existing knowledge.

### tRNA arrays

tRNA arrays were detected in isolates GM17 and OCU479 (Supplementary Table 6). tRNA array was discovered in some MAH isolates in the past study, and phylogenetic analysis based on nucleotide sequences of tRNA array showed that tRNA array of MAH was classified into a specific group [35]. To confirm tRNA arrays in this study as authentic tRNA arrays, phylogenetic analysis was performed. Our tRNA arrays were classified into the group 3, as defined in a previous study (Fig. 3) [35].

**Figure 3.**
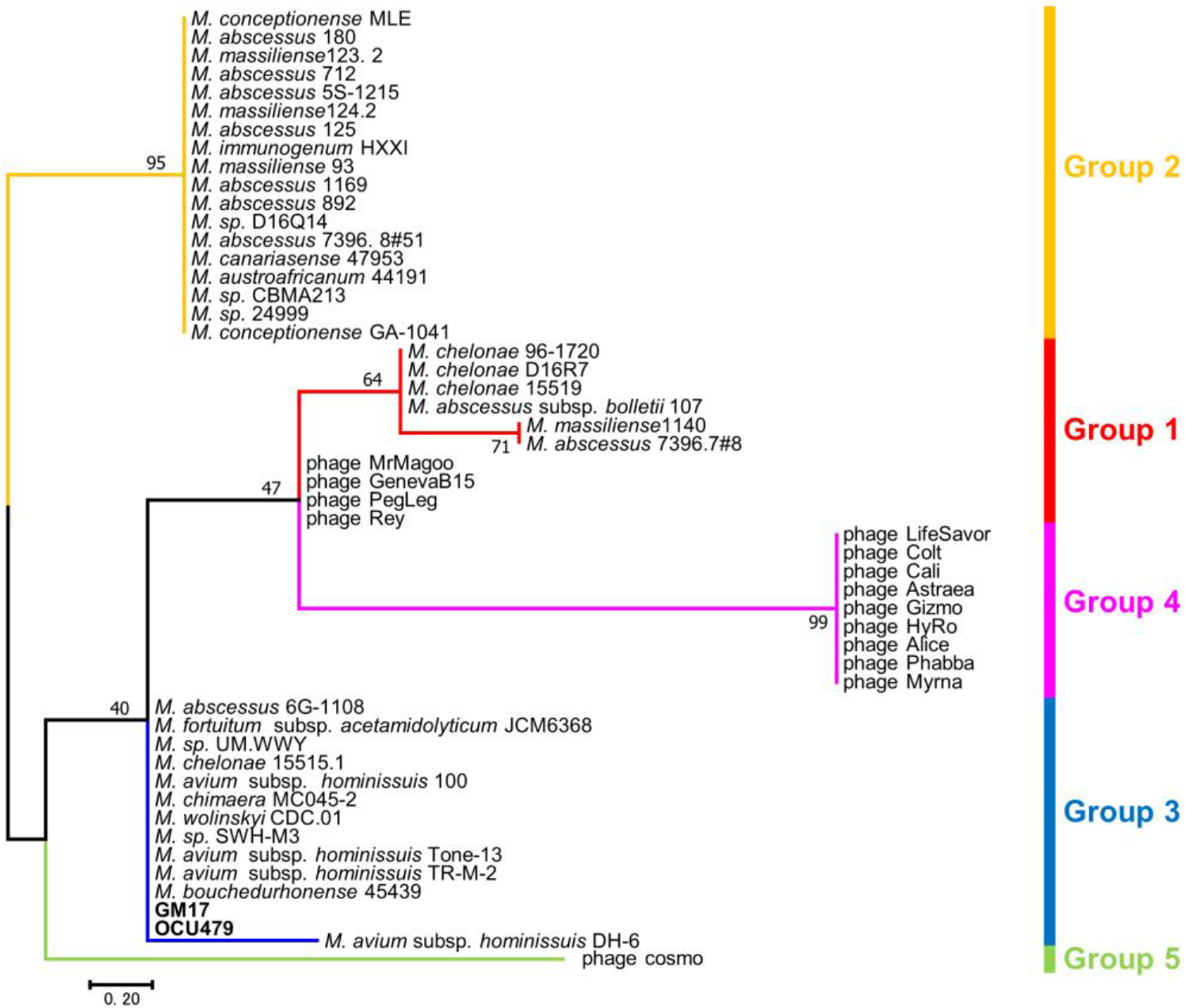
Phylogenetic tree based on the sequence of tRNA isotype located in tRNA array. Phylogenetic tree was generated by maximum likelihood method using MEGA 7.0. Two isolates (GM17 and OCU479 indicated in bold) were classified in Group 3. The bootstrap values were determined from 1,000 replications. The scale bar indicates genetic distances among strains.

## Re-use potential

MAH is known as one of the most critical *M. avium* subspecies causing non-tuberculosis mycobacterial infection in human and pigs. Pigs are suspected to be the most dominant host of MAH in animal and a potential source of infection for human [7, 8, 9, 10]. However, the study about relationship with human and pig MAH isolates based on genome is limited [11, 12]. Our study provides 30 draft genome sequences of MAH isolated from pigs. We believe that these data will be useful for genome-based epidemiological studies to evaluate the importance of pigs as a source of infection. In addition, we provide molecular identification of defense systems, tRNA arrays, virulence factors and drug resistance genes. These data are expected to be used in future research of MAH classification, pathogenicity, and identification of antimicrobial drug targets. Principally, our draft genomes were derived from both cases of systemic and lymph node limited infection of MAH. Thus, the provided virulence factors can be included in the important candidate genes associated with systemic infection of pigs.

## Data Availability

The summary information of draft genomes of the 30 MAH isolates are shown in Table 1. The genome sizes ranged to from approximately 4.8Mbps to 5.6Mbps. GC content was from 68.77% to 69.26%. All genome sequences have been deposited in GenBank under accession numbers VRUQ00000000, WEGO00000000 to WEGZ00000000 and WEHA00000000 to WEHQ00000000, and SRA under accession numbers SRR13521605, SRR13556487 to SRR13556515.

## Declarations

### List of abbreviations

NTM: non-tuberculous mycobacteria
MAC: *Mycobacterium avium* complex
MAA: *M. avium* subsp. *avium*
MAP: *M. avium* subsp. *paratuberculosis*
MAS: *M. avium* subsp. *silvaticum*
MAH: *Mycobacterium avium* subsp. *hominissuis*
MTB: *Mycobacterium tuberculosis*
RM: restriction modification
CRISPR: clustered regularly interspaced short palindromic repeat
BLAST: Basic Local Alignment Search Tool
MEGA: Molecular Evolutionary Genetics Analysis
NCBI: National Center for Biotechnology Information
PGAP: Prokaryotic Genome Annotation Pipeline
SNP: single nucleotide polymorphism

## Consent for publication

Not applicable.

## Author’s contributions

T.K, K.O and H.Y wrote the manuscript. K.M, A.H, S.S and K.S collected samples. K.O, J.O.O, S.S and K.S performed laboratory works. T.K, K.O, A.O, H.Y, J.O.O, T.Ito and M.K conducted computational analysis. Y.N, T.A, T.Y, H.F, T.W, S.Y, K.A designed methods. M.A, A.D.B, K.O, N.Y, T.Iwamoto and F.M designed whole research and advised on the interpretation of the study’s findings. All authors reviewed the manuscript.

## Acknowledgements

We thank the member of Gifu central hygiene service center and Toyama meat inspection center for sampling. Computational resources were partly provided by the Data Integration and Analysis Facility, National Institute for Basic Biology, Japan.

## Tables

**Supplementary Table 1. Isolates information and molecular characteristics of 30 MAH in this study**. a: Detected IS was 1213bp and shared 83% identity with IS*900*.

**Supplementary Table 2. Restriction modification system detected in 30 MAH isolates in this study**. ^*1^: These genes include the function of restriction enzyme/methyltransferase. ^*2^: These genes could be orphan methyltransferase. Yellow background: putative genes.

**Supplementary Table 3. Detected CRISPR-Cas systems in MAH GM44**.

**Supplementary Table 4. Virulence factors detected in 30 MAH isolates in this study**.

**Supplementary Table 5. Drug resistance genes detected in 30 MAH isolates in this study**.

**Supplementary Table 6. The information about tRNA array detected in MAH isolates GM17 and OCU479**.

## References

1. Daley CL. Mycobacterium avium complex disease. Microbiol Spectr. 2017;5.

2. Uchiya K, Takahashi H, Nakagawa T, Yagi T, Moriyama M, Inagaki T, et al. Characterization of a novel plasmid, pMAH135, from Mycobacterium avium subsp. hominissuis. PLoS One. 2015;10:e0117797.

3. Uchiya KI, Asahi S, Futamura K, Hamaura H, Nakagawa T, Nikai T, et al. Antibiotic susceptibility and genotyping of Mycobacterium avium strains that cause pulmonary and disseminated infection. Antimicrob. Agents Chemother. 2018;62:e02035–17.

4. Agdestein A, Johansen TB, Polaček V, Lium B, Holstad G, Vidanović D, et al. Investigation of an outbreak of mycobacteriosis in pigs. BMC Vet Res. 2011;7;:63.

5. Agdestein A, Johansen TB, Kolbjørnsen Ø, Jørgensen A, Djønne B, Olsen I, et al. A comparative study of Mycobacterium avium subsp. avium and Mycobacterium avium subsp. hominissuis in experimentally infected pigs. BMC Vet Res. 2012;8:11.

6. Hibiya K, Kasumi Y, Sugawara I Fujita J. Histopathological classification of systemic Mycobacterium avium complex infections in slaughtered domestic pigs. Comp Immunol Microbiol Infect Dis. 2008;31:347–66.

7. Agdestein A, Olsen I, Jørgensen A, Djønne B, Johansen TB. Novel insights into transmission routes of Mycobacterium avium in pigs and possible implications for human health. Vet Res. 2014;45:46.

8. Johansen TB, Olsen I, Jensen MR, Dahle UR, Holstad G, Djønne B. New probes used for IS1245 and IS1311 restriction fragment length polymorphism of Mycobacterium avium subsp. avium and Mycobacterium avium subsp. hominissuis isolates of human and animal origin in Norway. BMC Microbiol. 2007;7:14..

9. Klanicova B, Slana I, Vondruskova H, Kaevska M, Pavlik I. Real-time quantitative PCR detection of Mycobacterium avium subspecies in meat products. J Food Prot. 2011;74:636–40.

10. Slana I, Kaevska M, Kralik P, Horvathova A, Pavlik I. Distribution of Mycobacterium avium subsp. avium and M. a. hominissuis in artificially infected pigs studied by culture and IS901 and IS1245 quantitative real time PCR. Vet Microbiol. 2010;144:437–43.

11. Yano H, Iwamoto T, Nishiuchi Y, Nakajima C, Starkova DA, Mokrousov I, et al. Population structure and local adaptation of MAC lung disease agent Mycobacterium avium subsp. hominissuis. Genome Biol Evol. 2017;9:2403–17.

12. Yano H, Suzuki H, Maruyama F, Iwamoto T. The recombination-cold region as an epidemiological marker of recombinogenic opportunistic pathogen Mycobacterium avium. BMC Genomics. 2019;20:752.

13. Dragset MS, Ioerger TR, Loevenich M, Haug M, Sivakumar N, Marstad A, et al. Global assessment of Mycobacterium avium subsp. hominissuis genetic requirement for growth and virulence. mSystems. 2019;4:e00402–19.

14. Bruffaerts N, Vluggen C, Roupie V, Duytschaever L, Van den Poel C, Denoël J, et al. Virulence and immunogenicity of genetically defined human and porcine isolates of M. avium subsp. hominissuis in an experimental mouse infection. PLoS One. 2017;12:e0171895.

15. dx.doi.org/10.17504/protocols.io.bujenuje

16. dx.doi.org/10.17504/protocols.io.bupvnvn6

17. Chen ZH, Butler WR, Baumstark BR. Ahearn DG. Identification and differentiation of Mycobacterium avium and M. intracellulare by PCR. J Clin Microbiol. 1996;34:1267–9.

18. Kim BJ, Lee SH, Lyu MA, Kim SJ, Bai GH, Chae GT, et al. Identification of mycobacterial species by comparative sequence analysis of the RNA polymerase gene (rpoB). J Clin Microbiol. 1999;37:1714–20.

19. Turenne CY, Semret M, Cousins, DV, Collins DM, Behr MA. Sequencing of hsp65 distinguishes among subsets of the Mycobacterium avium complex. J. Clin. Microbiol. 2006;44:433–40.

20. Ahrens P, Giese SB., Klausen J, Inglis NF. Two markers, IS901-IS902 and p40, identified by PCR and by using monoclonal antibodies in Mycobacterium avium strains. J Clin Microbiol.1995;33:1049–53.

21. Kunze ZM, Wall S, Appelberg R, Silva MT, Portaels F, McFadden JJ. IS901, a new member of a widespread class of atypical insertion sequences, is associated with pathogenicity in Mycobacterium avium. Mol Microbiol. 1991;5:2265–72.

22. Sanderson JD, Moss MT, Tizard ML, Hermon-Taylor J. Mycobacterium paratuberculosis DNA in Crohn’s disease tissue. Gut. 1992;33:890–6.

23. Siguier P, Perochon J, Lestrade L, Mahillon J. Chandler M. ISfinder: the reference center for bacterial insertion sequences. Nucleic Acids Res. 2006;34:D32–6.

24. Nurk S, Bankevich A, Antipov D, Gurevich AA, Korobeynikov A, Lapidus A, et al. Assembling single-cell genomes and mini-metagenomes from chimeric MDA products. J Comput Biol. 2013;20:714–37.

25. Walker BJ, Abeel T, Shea T, Priest M, Abouelliel A, Sakthikumar S, et al. Pilon: an integrated tool for comprehensive microbial variant detection and genome assembly improvement. PLoS One. 2014;9:e112963.

26. Wick RR, Judd LM, Gorrie CL, Holt KE. Unicycler: Resolving bacterial genome assemblies from short and long sequencing reads. PLoS Comput Biol. 2017;13:e1005595.

27. Parks DH. Imelfort M, Skennerton C T, Hugenholtz P. Tyson GW. CheckM: assessing the quality of microbial genomes recovered from isolates, single cells, and metagenomes. Genome Res. 2015;25:1043–55.

28. Menzel P, Ng KL, Krogh A. Fast and sensitive taxonomic classification for metagenomics with Kaiju. Nat Commun. 2016;7:11257.

29. Eren AM, Esen ÖC, Quince C, Vineis JH, Morrison HG, Sogin ML, Anvi’o: an advanced analysis and visualization platform for ‘omics data. PeerJ. 2015;3:e1319.

30. Tatusova T, DiCuccio M, Badretdin A, Chetvernin V, Nawrocki EP, Zaslavsky L, et al. NCBI prokaryotic genome annotation pipeline. Nucleic Acids. Res. 2016;44:6614–24.

31. Roer L, Hendriksen RS, Leekitcharoenphon P, Lukjancenko O, Kaas RS, Hasman H, et al. Is the Evolution of Salmonella enterica subsp. enterica Linked to Restriction-Modification Systems? mSystems. 2016;1:e00009–16.

32. Couvin D, Bernheim A, Toffano-Nioche C, Touchon M, Michalik J, Néron B, et al. CRISPRCasFinder, an update of CRISRFinder, includes a portable version, enhanced performance and integrates search for Cas proteins. Nucleic Acids Res. 2018;46:W246–51.

33. Grissa I, Vergnaud G, Pourcel C. CRISPRFinder: a web tool to identify clustered regularly interspaced short palindromic repeats. Nucleic Acids Res. 2007;35:W52–7.

34. Lowe TM, Chan PP. tRNAscan-SE On-line: integrating search and context for analysis of transfer RNA genes. Nucleic Acids Res. 2016;44:W54–7.

35. Morgado SM, Vicente ACP. Beyond the Limits: tRNA array units in Mycobacterium genomes. Front Microbiol. 208;9:1042.

36. Liu B, Zheng D, Jin Q, Chen L, Yang J. VFDB 2019: a comparative pathogenomic platform with an interactive web interface. Nucleic Acids Res. 2019;47:D687–92.

37. Alcock BP, Raphenya AR, Lau TTY, Tsang KK, Bouchard M, Edalatmand A, et al. CARD 2020: antibiotic resistome surveillance with the comprehensive antibiotic resistance database. Nucleic Acids Res. 2020;48:D517–25.

38. Ichikawa K, Yagi T, Moriyama M, Inagaki T, Nakagawa T, Uchiya KI, et al. Characterization of Mycobacterium avium clinical isolates in Japan using subspecies-specific insertion sequences, and identification of a new insertion sequence, ISMav6. J Med Microbiol. 2009;58:945–50.

39. Mijs W, de Haas P, Rossau R, Van der Laan T, Rigouts L, Portaels F, et al. Molecular evidence to support a proposal to reserve the designation Mycobacterium avium subsp. avium for bird-type isolates and ‘M. avium subsp. hominissuis’ for the human/porcine type of M. avium. Int J Syst Evol Microbiol.2002;52:1505–18.

40. Hibiya K, Kazumi Y, Nishiuchi Y, Sugawara I, Miyagi K, Oda Y, et al. Descriptive analysis of the prevalence and the molecular epidemiology of Mycobacterium avium complex-infected pigs that were slaughtered on the main island of Okinawa. Comp Immunol Microbiol Infect Dis. 2010;33:401–21.

41. Matern WM, Bader JS, Karakousis PC. Genome analysis of Mycobacterium avium subspecies hominissuis strain 109. Sci Data. 2018;5:180277.

42. Uchiya K, Takahashi H, Yagi T, Moriyama M, Inagaki T, Ichikawa K, et al. Comparative genome analysis of Mycobacterium avium revealed genetic diversity in strains that cause pulmonary and disseminated disease. PLoS One. 2013;8:e71831.

43. Gorla P, Plocinska R, Sarva K, Satsangi AT, Pandeeti E, Donnelly R, et al. MtrA response regulator controls cell division and cell wall metabolism and affects susceptibility of Mycobacteria to the first line antituberculosis drugs. Front Microbiol. 2018;9:2839.

44. Newell KV, Thomas DP, Brekasis D, Paget MS. The RNA polymerase-binding protein RbpA confers basal levels of rifampicin resistance on Streptomyces coelicolor. Mol Microbiol. 2006;60:687–96.

