## Supplementary Tables for "Genomic features of *Mycobacterium avium* subsp. *hominissuis* isolated from pigs in Japan": Supplementary_Table 1.pdf

| Isolate | Farm | Arera | Collection date | Source | Lesion distribution | IS*900* | IS*901* | IS*902* | IS*1245* | IS*1311* | IS*1613* | *hsp* code |
|---|---|---|---|---|---|---|---|---|---|---|---|---|
| GM5 | Gifu A | Tokai | 15-Jul-2015 | mesenteric lymph nodes | Lymphadenitis | - | - | - | + | + | - | 2 |
| GM10 | Gifu A | Tokai | 29-Jul-2015 | mesenteric lymph nodes | Lymphadenitis | - | - | - | - | + | - | N |
| GM12 | Shiga A | Tokai | 05-Aug-2015 | mesenteric lymph nodes | Lymphadenitis | - | - | - | - | + | - | N |
| GM16 | Aichi A | Tokai | 01-Sep-2015 | mesenteric lymph nodes | Lymphadenitis | - | - | - | + | + | - | N |
| GM17 | Aichi B | Tokai | 01-Sep-2015 | mesenteric lymph nodes | Lymphadenitis | - | - | - | + | + | - | 1 |
| GM21 | Aichi B | Tokai | 04-Sep-2015 | mesenteric lymph nodes | Lymphadenitis | - | - | - | - | + | - | N |
| GM32 | Gifu A | Tokai | 05-Oct-2015 | mesenteric lymph nodes | Lymphadenitis | - | - | - | - | + | - | N |
| GM44 | Gifu B | Tokai | 24-Nov-2015 | mesenteric lymph nodes | Lymphadenitis | - | - | - | + | + | - | 1 |
| OCU467 | Toyama A1 | Hokuriku | 25-Aug-1999 | mesenteric lymph nodes | Systemic | - | - | - | + | + | - | 1 |
| OCU468 | Toyama A1 | Hokuriku | 01-Dec-1999 | mesenteric lymph nodes | Systemic | - | - | - | - | + | - | 1 |
| OCU469 | Ishikawa B | Hokuriku | 24-Dec-1999 | mesenteric lymph nodes | Systemic | - | - | - | + | + | - | 2 |
| OCU470 | Toyama A1 | Hokuriku | 24-Dec-1999 | mesenteric lymph nodes | Systemic | - | - | - | - | + | - | 1 |
| OCU471 | Ishikawa B | Hokuriku | 28-Jan-2000 | mesenteric lymph nodes | Systemic | - | - | - | + | + | - | 1 |
| OCU472 | Gifu B | Tokai | 04-Feb-2000 | mesenteric lymph nodes | Systemic | - | - | - | - | + | - | 1 |
| OCU473 | Ishikawa B | Hokuriku | 08-Feb-2000 | mesenteric lymph nodes | Systemic | - | - | - | + | - | - | 1 |
| OCU474 | Ishikawa B | Hokuriku | 22-Dec-2000 | mesenteric lymph nodes | Systemic | - | - | - | + | + | - | 1 |
| OCU475 | Toyama D | Hokuriku | 05-Jan-2001 | mesenteric lymph nodes | Systemic | - | - | - | - | + | - | 1 |
| OCU476 | Toyama D | Hokuriku | 11-Jan-2001 | mesenteric lymph nodes | Systemic | - | - | - | - | + | - | 1 |
| OCU477 | Toyama D | Hokuriku | 11-Jan-2001 | mesenteric lymph nodes | Systemic | - | - | - | + | + | - | 1 |
| OCU478 | Toyama A1 | Hokuriku | 18-Jan-2001 | mesenteric lymph nodes | Systemic | - | - | - | + | + | - | 1 |
| OCU479 | Ishikawa B | Hokuriku | 23-Jan-2001 | mesenteric lymph nodes | Systemic | - | - | - | - | + | - | 1 |
| OCU480 | Ishikawa B | Hokuriku | 26-Jan-2001 | mesenteric lymph nodes | Systemic | - | - | - | + | + | - | 1 |
| OCU481 | Toyama D | Hokuriku | 06-Feb-2001 | mesenteric lymph nodes | Systemic | - | - | - | + | + | +a | 1 |
| OCU482 | Ishikawa B | Hokuriku | 09-Feb-2001 | mesenteric lymph nodes | Systemic | - | - | - | + | + | +a | 1 |
| OCU483 | Ishikawa B | Hokuriku | 02-Dec-2004 | mesenteric lymph nodes | Lymphadenitis | - | - | - | + | + | - | 2 |
| OCU484 | Toyama A2 | Hokuriku | 02-Dec-2004 | mesenteric lymph nodes | Lymphadenitis | - | - | - | + | + | - | 3 |
| OCU485 | Toyama A3 | Hokuriku | 03-Dec-2004 | mesenteric lymph nodes | Lymphadenitis | - | - | - | + | + | - | 1 |
| OCU486 | Toyama A1 | Hokuriku | 06-Dec-2004 | mesenteric lymph nodes | Systemic | - | - | - | + | + | - | 1 |
| Toy194 | Toyama E | Hokuriku | 26-Mar-2018 | liver | Systemic | - | - | - | + | + | - | 1 |
| Toy195 | Toyama E | Hokuriku | 26-Mar-2018 | mandibular lymph nodes | Systemic | - | - | - | + | + | - | 1 |
