## Supplementary Tables for "Genomic features of *Mycobacterium avium* subsp. *hominissuis* isolated from pigs in Japan": Supplementary_Table 2_1st response.pdf

| | | | 1. Restriction enzyme | | | | | 2. Restriction enzyme | | | | | 1. Methyltransferase | | | | | 2. Methyltransferase | | | | | 1. specificity subunit | | | | | 2. specificity subunit | | | | |
|---|---|---|---|---|---|---|---|---|---|---|---|---|---|---|---|---|---|---|---|---|---|---|---|---|---|---|---|---|---|---|---|---|
| Isolate | Type | Contig | Gene | Identity (%) | Query/HSP | Position in contig | Recognition sequence | Gene | Identity (%) | Query/HSP | Position in contig | Recognition sequence | Gene | Identity (%) | Query/HSP | Position in contig | Recognition sequence | Gene | Identity (%) | Query/HSP | Position in contig | Recognition sequence | Gene | Identity (%) | Query/HSP | Position in contig | Recognition sequence | Gene | Identity (%) | Query/HSP | Position in contig | Recognition sequence |
| GM5 | putative Type I | contig_31_length_38995_depth_0.97x | Mav135ORF3057P | 91.59 | 3210 / 3210 | 24423..25979 | Unknown | - | - | - | - | - | M.Mtu30073II | 84.85 | 1662/1558 | 28322..31531 | Unknown | - | - | - | - | - | - | - | - | - | - | - | - | - | - | - |
| | putative Type IIG | contig_170_length_5550_depth_0.60x | Mav104ORF4226P[*1] | 100 | 1581 / 1581 | 3068..4648 | CTGGAG | - | - | - | - | - | M.MkaDI | 81.51 | 1578/557 | 3267..3822 | CTGGAG | Mav104ORF4226P | 100 | 1581 / 1581 | 3068..4648 | Unknown | - | - | - | - | - | - | - | - | - | - |
| GM10 | putative Type II | contig_82_length_19162_depth_0.74x | - | - | - | - | - | - | - | - | - | - | M.MkaDI | 81.54 | 1578/558 | 11824..12379 | CTGGAG | M.Mav93ORF1950P | 99.68 | 1581 / 1581 | - | Unknown | - | - | - | - | - | - | - | - | - | - |
| GM12 | putative Type II | contig_111_length_14167_depth_0.70x | - | - | - | - | - | - | - | - | - | - | M.MkaDI | 81.54 | 1578/558 | 6814..7369 | CTGGAG | M.Mav93ORF1950P | 99.68 | 1581 / 1581 | 5988..7568 | Unknown | - | - | - | - | - | - | - | - | - | - |
| GM16 | putative Type I | contig_4_length_59942_depth_1.10x | Mav135ORF3057P | 91.59 | 3210 / 3210 | 12827..16036 | Unknown | - | - | - | - | - | M.Mtu30073II | 84.85 | 1662/1558 | 18379..19935 | Unknown | - | - | - | - | - | - | - | - | - | - | - | - | - | - | - |
| | putative Type I | contig_194_length_8291_depth_1.07x | Mav2285ORF23555P | 100 | 2364 / 2364 | 4472..6835 | Unknown | - | - | - | - | - | M.Mav104ORF5027P | 100 | 1488 / 1488 | 1453..2940 | Unknown | - | - | - | - | - | S2.Mav2285ORF23555P | 100 | 447 / 447 | 383..829 | Unknown | S1.Mav2285ORF23555P | 100 | 432 / 432 | 1004..1435 | Unknown |
| | putative Type IIG | contig_247_length_4646_depth_0.49x | Mav104ORF4226P[*1] | 100 | 1581 / 1581 | 2169..3749 | Unknown | - | - | - | - | - | M.MkaDI | 81.51 | 1578/557 | 2368..2923 | CTGGAG | - | - | - | - | - | - | - | - | - | - | - | - | - | - | - |
| GM17 | putative Type I | contig_66_length_25175_depth_1.12x | MavHMCO2P(MavHMCO2BP) | 99.13 | 1381 / 1380 | 4177..5553 | Unknown | - | - | - | - | - | M.MavHMCO2P(M.MavHMCO2AP) | 99.08 | 760 / 789 | 5550..6306 | Unknown | M.MavHMCO2P(M.MavHMCO2BP) | 98.38 | 495 / 495 | 6306..6797 | Unknown | - | - | - | - | - | - | - | - | - | - |
| | putative Type IIG | contig_157_length_10215_depth_0.60x | Mav104ORF4226P[*1] | 99.18 | 1581 / 1581 | 2652..4232 | Unknown | - | - | - | - | - | M.MkaDI | 81.72 | 1578/558 | 2851..3406 | CTGGAG | - | - | - | - | - | - | - | - | - | - | - | - | - | - | - |
| GM21 | putative Type II | contig_75_length_19160_depth_0.74x | - | - | - | - | - | - | - | - | - | - | M.MkaDI | 81.54 | 1578/558 | 6812..7367 | CTGGAG | M.Mav93ORF1950P | 99.68 | 1581 / 1581 | 5986..7566 | Unknown | - | - | - | - | - | - | - | - | - | - |
| GM32 | putative Type II | contig_65_length_23771_depth_0.89x | - | - | - | - | - | - | - | - | - | - | M.MkaDI | 81.54 | 1578/558 | 20846..21401 | CTGGAG | M.Mav93ORF1950P | 99.68 | 1581 / 1581 | 20020..21600 | Unknown | - | - | - | - | - | - | - | - | - | - |
| GM44 | putative Type I | contig_189_length_8266_depth_1.21x | M.Mav104ORF5027P | 100 | 1488 / 1488 | 1451..2938 | Unknown | Mav2285ORF23555P | 100 | 2364 / 2364 | 4470..6833 | Unknown | - | - | - | - | - | - | - | - | - | - | S2.Mav2285ORF23555P | 99.78 | 447 / 447 | 381..827 | Unknown | S1.Mav2285ORF23555P | 100 | 432 / 432 | 1002..1433 | Unknown |
| | putative Type I | contig_183_length_8434_depth_1.08x | MavHMCO2P(MavHMCO2BP) | 99.13 | 1381 / 1380 | 6620..7996 | Unknown | - | - | - | - | - | M.MavHMCO2P(M.MavHMCO2BP) | 98.38 | 495 / 495 | 5376..5867 | Unknown | M.MavHMCO2P(M.MavHMCO2AP) | 99.08 | 760 / 789 | 5867..6623 | Unknown | - | - | - | - | - | - | - | - | - | - |
| | putative Type IIG | contig_155_length_10660_depth_0.69x | Mav104ORF4226P[*1] | 100 | 1581 / 1581 | 3094..4674 | Unknown | - | - | - | - | - | M.MkaDI | 81.51 | 1578/557 | 3293..3848 | CTGGAG | - | - | - | - | - | - | - | - | - | - | - | - | - | - | - |
| | putative Type III | contig_5_length_77846_depth_1.06x | Mav104ORF275P | 100 | 2592 / 2592 | 73309..75900 | Unknown | - | - | - | - | - | M.Mav104ORF275P | 100 | 1877 / 1989 | 75900..77776 | Unknown | - | - | - | - | - | - | - | - | - | - | - | - | - | - | - |
| OCU467 | putative Type II | contig_8_length_269030_depth_0.98x | Mav4293ORF19350P | 100 | 780 / 780 | 173030..173809 | GATC | - | - | - | - | - | M.MgoX7BORFAP | 100 | 840 / 840 | 173915..174754 | RGATCY | - | - | - | - | - | - | - | - | - | - | - | - | - | - | - |
| | putative Type IIG | contig_12_length_187613_depth_0.99x | Mav104ORF4226P[*1] | 100 | 1581 / 1581 | - | Unknown | - | - | - | - | - | M.MkaDI | 81.51 | 1578/557 | 89016..89571 | CTGGAG | - | - | - | - | - | - | - | - | - | - | - | - | - | - | - |
| OCU468 | putative Type I | contig_55_length_14458_depth_1.01x | Mav2285ORF23555P | 100 | 2364 / 2364 | 7693..10056 | Unknown | - | - | - | - | - | M.Mav104ORF5027P | 100 | 1488 / 1488 | 11588..13075 | Unknown | - | - | - | - | - | S1.Mav2285ORF23555P | 100 | 432 / 432 | 13093..13524 | - | S2.Mav2285ORF23555P | 100 | 447 / 447 | 13699..14145 | Unknown |
| | putative Type I | contig_27_length_63452_depth_1.02x | Mav135ORF3057P | 91.59 | 3210 / 3210 | 53372..56581 | Unknown | - | - | - | - | - | - | - | - | - | - | - | - | - | - | - | - | - | - | - | - | - | - | - | - | - |
| | putative Type II | contig_52_length_15208_depth_1.02x | - | - | - | - | - | - | - | - | - | - | - | - | - | - | - | - | - | - | - | - | S.Meu1344ORF144P | 100 | 789 / 981 | 1..789 | Unknown | - | - | - | - | - |
| | putative Type IIG | contig_8_length_160272_depth_1.02x | Mav104ORF4226P[*1] | 100 | 1581 / 1581 | 64908..66488 | Unknown | - | - | - | - | - | M.MkaDI | 81.51 | 1578/557 | 65107..65662 | CTGGAG | - | - | - | - | - | - | - | - | - | - | - | - | - | - | - |
| OCU469 | putative Type I | contig_13_length_168214_depth_0.99x | Mav135ORF3057P | 91.59 | 3210 / 3210 | 53107..56316 | Unknown | - | - | - | - | - | M.Mtu30073II | 84.85 | 1662/1558 | 49208..50764 | Unknown | - | - | - | - | - | - | - | - | - | - | - | - | - | - | - |
| | putative Type IIG | contig_9_length_190329_depth_0.98x | Mav104ORF4226P[*1] | 100 | 1581 / 1581 | 91323..92903 | Unknown | - | - | - | - | - | M.MkaDI | 81.51 | 1578/557 | 91522..92077 | CTGGAG | - | - | - | - | - | - | - | - | - | - | - | - | - | - | - |
| OCU470 | putative Type I | contig_60_length_14457_depth_0.96x | Mav2285ORF23555P | 100 | 2364 / 2364 | 7692..10055 | Unknown | - | - | - | - | - | M.Mav104ORF5027P | 100 | 1488 / 1488 | 11587..13074 | Unknown | - | - | - | - | - | S1.Mav2285ORF23555P | 100 | 432 / 432 | 13092..13523 | Unknown | S2.Mav2285ORF23555P | 100 | 447 / 447 | 13698..14144 | Unknown |
| | putative Type I | contig_30_length_63452_depth_0.99x | Mav135ORF3057P | 91.59 | 3210 / 3210 | 6872..10081 | Unknown | - | - | - | - | - | - | - | - | - | - | - | - | - | - | - | - | - | - | - | - | - | - | - | - | - |
| | putative Type II | contig_52_length_16972_depth_0.91x | - | - | - | - | - | - | - | - | - | - | - | - | - | - | - | - | - | - | - | - | S.Meu1344ORF144P | 100 | 789 / 981 | 1..789 | Unknown | - | - | - | - | - |
| | putative Type IIG | contig_16_length_112060_depth_1.01x | Mav104ORF4226P[*1] | 99.94 | 1581 / 1581 | 64908..66488 | Unknown | - | - | - | - | - | M.MkaDI | 81.51 | 1578/557 | 65107..65662 | CTGGAG | - | - | - | - | - | - | - | - | - | - | - | - | - | - | - |
| OCU471 | putative Type II | contig_22_length_66407_depth_1.00x | Mav4293ORF19350P | 100 | 780 / 780 | 9594..10373 | Unknown | - | - | - | - | - | M.MgoX7BORFAP | 100 | 840 / 840 | 10479..11318 | Unknown | - | - | - | - | - | - | - | - | - | - | - | - | - | - | - |
| | putative Type IIG | contig_14_length_112000_depth_1.00x | Mav104ORF4226P[*1] | 100 | 1581 / 1581 | 45568..47148 | Unknown | - | - | - | - | - | M.MkaDI | 81.51 | 1578/557 | 46394..46949 | CTGGAG | - | - | - | - | - | - | - | - | - | - | - | - | - | - | - |
| OCU472 | putative Type I | contig_64_length_14457_depth_1.00x | Mav2285ORF23555P | 100 | 2364 / 2364 | 7692..10055 | Unknown | - | - | - | - | - | M.Mav104ORF5027P | 100 | 1488 / 1488 | 11587..13074 | Unknown | - | - | - | - | - | S1.Mav2285ORF23555P | 100 | 432 / 432 | 13092..13523 | Unknown | S2.Mav2285ORF23555P | 100 | 447 / 447 | 13698..14144 | Unknown |
| | putative Type I | contig_29_length_63452_depth_1.01x | Mav135ORF3057P | 91.59 | 3210 / 3210 | 6872..10081 | Unknown | - | - | - | - | - | - | - | - | - | - | - | - | - | - | - | - | - | - | - | - | - | - | - | - | - |
| | putative Type II | contig_55_length_19836_depth_0.98x | - | - | - | - | - | - | - | - | - | - | - | - | - | - | - | - | - | - | - | - | S.Meu1344ORF144P | 100 | 789 / 981 | 19048..19836 | Unknown | - | - | - | - | - |
| | putative Type IIG | contig_17_length_112060_depth_1.00x | Mav104ORF4226P[*1] | 100 | 1581 / 1581 | 64908..66488 | Unknown | - | - | - | - | - | M.MkaDI | 81.51 | 1578/557 | 65107..65662 | CTGGAG | - | - | - | - | - | - | - | - | - | - | - | - | - | - | - |
| OCU473 | putative Type I | contig_16_length_105441_depth_0.99x | Mav135ORF3057P | 91.59 | 3210 / 3210 | 6873..10082 | Unknown | - | - | - | - | - | M.Mtu30073II | 84.85 | 1662/1558 | 2974..4530 | CTGGAG | - | - | - | - | - | - | - | - | - | - | - | - | - | - | - |
| | putative Type I | contig_28_length_80296_depth_0.93x | Mav2285ORF23555P | 100 | 2364 / 2364 | 73530..75893 | Unknown | - | - | - | - | - | M.Mav104ORF5027P | 100 | 1488 / 1488 | 77425..78912 | Unknown | - | - | - | - | - | S1.Mav2285ORF23555P | 100 | 432 / 432 | 78930..79361 | Unknown | S2.Mav2285ORF23555P | 100 | 447 / 447 | 79536..79982 | Unknown |
| | putative Type II | contig_3_length_232359_depth_0.97x | - | - | - | - | - | - | - | - | - | - | M.Mky204ORFAP[*2] | 100 | 930 / 930 | 216974..217903 | Unknown | - | - | - | - | - | - | - | - | - | - | - | - | - | - | - |
| | putative Type IIG | contig_14_length_112028_depth_1.05x | Mav104ORF4226P[*1] | 100 | 1581 / 1581 | 64908..66488 | Unknown | - | - | - | - | - | M.MkaDI | 81.51 | 1578/557 | 65107..65662 | CTGGAG | - | - | - | - | - | - | - | - | - | - | - | - | - | - | - |
| OCU474 | putative Type II | contig_2_length_508797_depth_1.01x | Mav4293ORF19350P | 100 | 780 / 780 | 344287..345066 | GATC | - | - | - | - | - | M.MgoX7BORFAP | 100 | 840 / 840 | 345172..346011 | RGATCY | - | - | - | - | - | - | - | - | - | - | - | - | - | - | - |
| | putative Type IIG | contig_9_length_163863_depth_1.01x | Mav104ORF4226P[*1] | 100 | 1581 / 1581 | 97370..98950 | Unknown | - | - | - | - | - | M.MkaDI | 81.51 | 1578/557 | 98196..98751 | CTGGAG | - | - | - | - | - | - | - | - | - | - | - | - | - | - | - |
| OCU475 | putative Type I | contig_62_length_14457_depth_0.99x | Mav2285ORF23555P | 100 | 2364 / 2364 | 4403..6766 | Unknown | - | - | - | - | - | M.Mav104ORF5027P | 100 | 1488 / 1488 | 1384..2871 | Unknown | - | - | - | - | - | S2.Mav2285ORF23555P | 100 | 447 / 447 | 314..760 | Unknown | S1.Mav2285ORF23555P | 100 | 432 / 432 | 935..1366 | Unknown |
| | putative Type I | contig_31_length_63452_depth_1.00x | Mav135ORF3057P | 91.59 | 3210 / 3210 | 6872..10081 | Unknown | - | - | - | - | - | - | - | - | - | - | - | - | - | - | - | - | - | - | - | - | - | - | - | - | - |
| | putative Type II | contig_56_length_18783_depth_0.97x | - | - | - | - | - | - | - | - | - | - | - | - | - | - | - | - | - | - | - | - | S.Meu1344ORF144P | 100 | 789 / 981 | 1..789 | Unknown | - | - | - | - | - |
| | putative Type IIG | contig_16_length_112060_depth_1.01x | Mav104ORF4226P[*1] | 100 | 1581 / 1581 | 45573..47153 | Unknown | - | - | - | - | - | M.MkaDI | 81.51 | 1578/557 | 46399..46954 | CTGGAG | - | - | - | - | - | - | - | - | - | - | - | - | - | - | - |
| OCU476 | putative Type I | contig_57_length_14457_depth_1.04x | Mav2285ORF23555P | 100 | 2364 / 2364 | 4403..6766 | Unknown | - | - | - | - | - | M.Mav104ORF5027P | 100 | 1488 / 1488 | 1384..2871 | Unknown | - | - | - | - | - | S2.Mav2285ORF23555P | 100 | 447 / 447 | 314..760 | Unknown | S1.Mav2285ORF23555P | 100 | 432 / 432 | 935..1366 | Unknown |
| | putative Type I | contig_28_length_63452_depth_0.98x | Mav135ORF3057P | 91.59 | 3210 / 3210 | 53372..56581 | Unknown | - | - | - | - | - | M.Mtu30073II | 84.85 | 1662/1558 | 58924..60480 | Unknown | - | - | - | - | - | - | - | - | - | - | - | - | - | - | - |
| | putative Type II | contig_50_length_16972_depth_1.00x | - | - | - | - | - | - | - | - | - | - | - | - | - | - | - | - | - | - | - | - | S.Meu1344ORF144P | 100 | 789 / 981 | 16184..16972 | Unknown | - | - | - | - | - |
| | putative Type IIG | contig_16_length_112060_depth_1.01x | Mav104ORF4226P[*1] | 99.94 | 1581 / 1581 | 64908..66488 | Unknown | - | - | - | - | - | M.MkaDI | 81.51 | 1578/557 | 65107..65662 | CTGGAG | - | - | - | - | - | - | - | - | - | - | - | - | - | - | - |
| OCU477 | putative Type II | contig_2_length_398853_depth_0.98x | Mav4293ORF19350P | 100 | 780 / 780 | 163732..164511 | GATC | - | - | - | - | - | M.MgoX7BORFAP | 100 | 840 / 840 | 162787..163626 | RGATCY | - | - | - | - | - | - | - | - | - | - | - | - | - | - | - |
| | putative Type IIG | contig_1_length_491303_depth_1.00x | Mav104ORF4226P[*1] | 100 | 1581 / 1581 | 64858..66438 | Unknown | - | - | - | - | - | M.MkaDI | 81.51 | 1578/557 | 65057..65612 | CTGGAG | - | - | - | - | - | - | - | - | - | - | - | - | - | - | - |
| OCU478 | putative Type II | contig_3_length_415117_depth_1.01x | Mav4293ORF19350P | 100 | 780 / 780 | 241308..242087 | GATC | - | - | - | - | - | M.MgoX7BORFAP | 100 | 840 / 840 | 240363..241202 | RGATCY | - | - | - | - | - | - | - | - | - | - | - | - | - | - | - |
| | putative Type IIG | contig_11_length_187613_depth_1.01x | Mav104ORF4226P[*1] | 100 | 1581 / 1581 | 97217..98797 | Unknown | - | - | - | - | - | M.MkaDI | 81.51 | 1578/557 | 98043..98598 | CTGGAG | - | - | - | - | - | - | - | - | - | - | - | - | - | - | - |
| OCU479 | putative Type I | contig_66_length_14457_depth_1.02x | Mav2285ORF23555P | 100 | 2364 / 2364 | 4403..6766 | Unknown | - | - | - | - | - | M.Mav104ORF5027P | 100 | 1488 / 1488 | 1384..2871 | Unknown | - | - | - | - | - | S2.Mav2285ORF23555P | 100 | 447 / 447 | 314..760 | Unknown | S1.Mav2285ORF23555P | 100 | 432 / 432 | 935..1366 | Unknown |
| | putative Type I | contig_29_length_63452_depth_0.99x | Mav135ORF3057P | 91.59 | 3210 / 3210 | 6872..10081 | Unknown | - | - | - | - | - | M.Mtu30073II | 84.85 | 1662/1558 | 2973..4529 | Unknown | - | - | - | - | - | - | - | - | - | - | - | - | - | - | - |
| | putative Type II | contig_59_length_19836_depth_1.05x | - | - | - | - | - | - | - | - | - | - | - | - | - | - | - | - | - | - | - | - | S.Meu1344ORF144P | 100 | 789 / 981 | 1..789 | Unknown | - | - | - | - | - |
| | putative Type IIG | contig_15_length_112152_depth_0.99x | Mav104ORF4226P[*1] | 99.94 | 1581 / 1581 | 45665..47245 | Unknown | - | - | - | - | - | M.MkaDI | 81.51 | 1578/557 | 46491..47046 | CTGGAG | - | - | - | - | - | - | - | - | - | - | - | - | - | - | - |
| | putative Type II | contig_70_length_12793_depth_1.59x | - | - | - | - | - | - | - | - | - | - | M.Mky204ORFAP[*2] | 100 | 930 / 930 | 8298..9227 | Unknown | - | - | - | - | - | - | - | - | - | - | - | - | - | - | - |
| OCU480 | putative Type II | contig_4_length_286713_depth_1.00x | Mav4293ORF19350P | 100 | 780 / 780 | 56017..56796 | GATC | - | - | - | - | - | M.MgoX7BORFAP | 100 | 840 / 840 | 55072..55911 | RGATCY | - | - | - | - | - | - | - | - | - | - | - | - | - | - | - |
| | putative Type IIG | contig_13_length_112819_depth_1.00x | Mav104ORF4226P[*1] | 100 | 1581 / 1581 | 64949..66529 | Unknown | - | - | - | - | - | M.MkaDI | 81.51 | 1578/557 | 65148..65703 | CTGGAG | - | - | - | - | - | - | - | - | - | - | - | - | - | - | - |
| | putative Type II | contig_1_length_533568_depth_0.99x | Mav4293ORF19350P | 100 | 780 / 780 | 415423..416202 | GATC | - | - | - | - | - | M.MgoX7BORFAP | 100 | 840 / 840 | 416308..417147 | RGATCY | - | - | - | - | - | - | - | - | - | - | - | - | - | - | - |
| OCU481 | putative Type IIG | contig_10_length_163705_depth_1.00x | Mav104ORF4226P[*1] | 100 | 1581 / 1581 | 97217..98797 | Unknown | - | - | - | - | - | M.MkaDI | 81.51 | 1578/557 | 98043..98598 | CTGGAG | - | - | - | - | - | - | - | - | - | - | - | - | - | - | - |
| OCU482 | putative Type II | contig_1_length_533568_depth_1.00x | Mav4293ORF19350P | 100 | 780 / 780 | 415423..416202 | GATC | - | - | - | - | - | M.MgoX7BORFAP | 100 | 840 / 840 | 416308..417147 | RGATCY | - | - | - | - | - | - | - | - | - | - | - | - | - | - | - |
| | putative Type IIG | contig_11_length_163705_depth_1.00x | Mav104ORF4226P[*1] | 100 | 1581 / 1581 | 97217..98797 | Unknown | - | - | - | - | - | M.MkaDI | 81.51 | 1578/557 | 98043..98598 | CTGGAG | - | - | - | - | - | - | - | - | - | - | - | - | - | - | - |
| OCU483 | putative Type I | contig_5_length_300223_depth_1.01x | MavHMCO2P(MavHMCO2BP) | 99.13 | 1381 / 1380 | 291348..292724 | Unknown | - | - | - | - | - | M.MavHMCO2P(M.MavHMCO2AP) | 99.08 | 760 / 789 | 292721..293477 | Unknown | M.MavHMCO2P(M.MavHMCO2BP) | 98.38 | 495 / 495 | 293477..293968 | Unknown | - | - | - | - | - | - | - | - | - | - |
| | putative Type IIG | contig_12_length_139140_depth_1.00x | Mav104ORF4226P[*1] | 99.37 | 1581 / 1581 | 46260..47840 | Unknown | - | - | - | - | - | M.MkaDI | 81.72 | 1578/558 | 47086..47641 | CTGGAG | - | - | - | - | - | - | - | - | - | - | - | - | - | - | - |
| OCU484 | putative Type I | contig_58_length_14422_depth_1.04x | Mav2285ORF23555P | 100 | 2364 / 2364 | 7657..10020 | Unknown | - | - | - | - | - | M.Mav104ORF5027P | 100 | 1488 / 1488 | 11552..13039 | Unknown | - | - | - | - | - | S1.Mav2285ORF23555P | 100 | 432 / 432 | 13057..13488 | Unknown | S2.Mav2285ORF23555P | 100 | 447 / 447 | 13663..14109 | Unknown |
| | putative Type I | contig_17_length_105828_depth_0.97x | Mav135ORF3057P | 91.59 | 3210 / 3210 | 95748..98957 | Unknown | - | - | - | - | - | M.Mtu30073II | 84.85 | 1662/1558 | 101300..102856 | Unknown | - | - | - | - | - | - | - | - | - | - | - | - | - | - | - |
| | putative Type IIG | contig_8_length_168109_depth_0.98x | Mav104ORF4226P[*1] | 99.49 | 1581 / 1581 | 94785..96365 | - | - | - | - | - | - | M.MkaDI | 81.83 | 1578/545 | 95611..96153 | CTGGAG | - | - | - | - | - | - | - | - | - | - | - | - | - | - | - |
| OCU485 | putative Type IIG | contig_11_length_187613_depth_1.01x | Mav104ORF4226P[*1] | 100 | 1581 / 1581 | 88817..90397 | Unknown | - | - | - | - | - | M.MkaDI | 81.51 | 1578/557 | 89016..89571 | CTGGAG | - | - | - | - | - | - | - | - | - | - | - | - | - | - | - |
| | putative Type II | contig_6_length_269137_depth_1.00x | Mav4293ORF19350P | 100 | 780 / 780 | 95274..96053 | GATC | - | - | - | - | - | M.MgoX7BORFAP | 100 | 840 / 840 | 94329..95168 | RGATCY | - | - | - | - | - | - | - | - | - | - | - | - | - | - | - |
| OCU486 | putative Type II | contig_7_length_248156_depth_0.96x | Mav4293ORF19350P | 100 | 780 / 780 | 56049..56828 | GATC | - | - | - | - | - | M.MgoX7BORFAP | 100 | 840 / 840 | 55104..55943 | RGATCY | - | - | - | - | - | - | - | - | - | - | - | - | - | - | - |
| | putative Type IIG | contig_13_length_112689_depth_1.02x | Mav104ORF4226P[*1] | 100 | 1581 / 1581 | 64857..66437 | Unknown | - | - | - | - | - | M.MkaDI | 81.51 | 1578/557 | 65056..65611 | CTGGAG | - | - | - | - | - | - | - | - | - | - | - | - | - | - | - |
| Toy194 | putative Type IIG | contig_3_length_491289_depth_1.00x | Mav104ORF4226P[*1] | 100 | 1581 / 1581 | 424857..426437 | Unknown | - | - | - | - | - | M.MkaDI | 81.51 | 1578/557 | 425683..426238 | CTGGAG | - | - | - | - | - | - | - | - | - | - | - | - | - | - | - |
| Toy195 | putative Type IIG | contig_1_length_491290_depth_1.00x | Mav104ORF4226P[*1] | 100 | 1581 / 1581 | 64853..66433 | Unknown | - | - | - | - | - | M.MkaDI | 81.51 | 1578/557 | 65052..65607 | CTGGAG | - | - | - | - | - | - | - | - | - | - | - | - | - | - | - |
