## Supplementary Tables for "Genomic features of *Mycobacterium avium* subsp. *hominissuis* isolated from pigs in Japan": Supplementary_Table 3_1st response.pdf

| Strain Name | Evidence level | Number of spacers | Sequnce of spacers | CRISPR start position | CRISPR end position | CRISPR length | Consensus Repeat | Repeat Length |
|---|---|---|---|---|---|---|---|---|
| GM44 | 4 | 12 | 1: ACCGGTCGGTCACTGCGGTGGTGTCCTGTGCATGCTCC<br>2: ACCTCCCAGGCGGACGCAGTGCCAGGGATGGCGAGTA<br>3: ACCCGAGGCCGTCGCGGAGGCCTTGACCGACCCCGATA<br>4: ACCGCGCACCTCAGCTGCTGTGCTGCGTGAGCGCGTCATA<br>5: ACCCCTGCACCAGTCGATCCACTGCGACGTGCGCAGCA<br>6: ACCCATCCCAGGTCAGGAAGTCTGCTCCCCGCGTAAGA<br>7: ACCGGGCCTGTTGCTCATCGGCCCGCCGCGCTCGGGCA<br>8: ACCGCCGATACCGGGCTTGGCATCCGTGCCGTACTGC<br>9: CCCCGTGCCCGGTGGAGGAACCACCTCTCCCCCCACA<br>10: ACCGGCCGCAGAGGAGGCCGTCACCGCGGCGAAGACC<br>11: ACCCCTCCGATCCAGGTACCGCGTCCGGAAGATGTGGCC<br>12: CCCCCCCCGTCTGCAGCGCAACGGTTCCTACTGCACCTCC | 4089 | 4860 | 771 | TGCTCCCCGCGCAAGCGGGGATGAACC | 27 |
