## Supplementary Tables for "Genomic features of *Mycobacterium avium* subsp. *hominissuis* isolated from pigs in Japan": Supplementary_Table 4_1st response.pdf

| | Amino acid and purine metabolism | | | | | | | | | | | Antiapoptosis factor | Catabolism of chol | |
|---|---|---|---|---|---|---|---|---|---|---|---|---|---|---|
| | Glutamine synthesis | Leucine synthesis | Lysine synthesis | Proline synthesis | Purine synthesis | Tryptophan synthesis | Nitrate reductase | | | | Nitrate/nitrite transporter | NuoG | Cyp125 | FadE28 |
| Isolates | glnA1 | leuD | lysA | proC | purC | trpD | narG | narH | narI | narJ | narK2 | nuoG | cyp125 | fadE28 |
| GM5 | + | + | + | + | + | + | + | + | + | + | - | - | + | + |
| GM10 | + | + | + | + | + | + | + | + | + | + | - | + | + | + |
| GM12 | + | + | + | + | + | - | + | + | + | + | - | + | + | + |
| GM16 | + | + | + | + | + | + | + | + | + | + | - | - | + | + |
| GM17 | + | + | + | + | + | - | + | + | + | + | - | + | + | + |
| GM21 | + | + | + | + | + | - | + | + | + | + | - | - | + | + |
| GM32 | + | + | + | + | + | - | + | + | + | + | - | - | + | + |
| GM44 | + | + | + | + | + | - | + | + | + | + | - | + | + | + |
| OCU467 | + | + | + | + | + | - | + | + | + | + | - | + | + | + |
| OCU468 | + | + | + | + | + | - | + | + | + | + | - | + | + | + |
| OCU469 | + | + | + | + | + | - | + | + | + | + | - | + | + | + |
| OCU470 | + | + | + | + | + | - | + | + | + | + | - | + | + | + |
| OCU471 | + | + | + | + | + | - | + | + | + | + | - | + | + | + |
| OCU472 | + | + | + | + | + | - | + | + | + | + | - | + | + | + |
| OCU473 | + | + | + | + | + | - | + | + | + | + | - | + | + | + |
| OCU474 | + | + | + | + | + | - | + | + | + | + | - | + | + | + |
| OCU475 | + | + | + | + | + | - | + | + | + | + | - | + | + | + |
| OCU476 | + | + | + | + | + | - | + | + | + | + | - | + | + | + |
| OCU477 | + | + | + | + | + | - | + | + | + | + | - | + | + | + |
| OCU478 | + | + | + | + | + | - | + | + | + | + | - | + | + | + |
| OCU479 | + | + | + | + | + | - | + | + | + | + | - | + | + | + |
| OCU480 | + | + | + | + | + | - | + | + | + | + | - | + | + | + |
| OCU481 | + | + | + | + | + | - | + | + | + | + | - | + | + | + |
| OCU482 | + | + | + | + | + | - | + | + | + | + | - | + | + | + |
| OCU483 | + | + | + | + | + | - | + | + | + | + | - | + | + | + |
| OCU484 | + | + | + | + | + | - | + | + | + | + | - | + | + | + |
| OCU485 | + | + | + | + | + | - | + | + | + | + | - | + | + | + |
| OCU486 | + | + | + | + | + | - | + | + | + | + | - | + | + | + |
| Toy194 | + | + | + | + | + | - | + | + | + | + | + | + | + | + |
| Toy195 | + | + | + | + | + | - | + | + | + | + | - | + | + | + |
| Total Number of isolates | 30 | 30 | 30 | 30 | 30 | 3 | 30 | 30 | 30 | 30 | 1 | 26 | 30 | 30 |

| esterol | | | | | | | | | | | | | | | | | | | |
|---|---|---|---|---|---|---|---|---|---|---|---|---|---|---|---|---|---|---|---|
| FadE29 | Carboxylesterase | Exported repetitive protein | GPL locus | | | | | | | | | | | | | | | | |
| fadE29 | caeA | erp | Undetermined | fad23 | fadE5 | gap-like | gap | gtf1 | gtf2 | gtf3 | mbtH | mmpL10 | mmpL4a | mmpL4b | mmpS4 | mps1 | mps2 | papA3 | pe |
| + | + | - | + | + | + | + | + | + | + | - | + | + | + | + | + | + | + | + | + |
| + | + | - | + | + | + | + | + | + | + | - | + | + | + | + | + | + | + | + | + |
| + | + | - | + | + | + | + | + | + | + | - | + | + | + | + | + | + | + | + | + |
| + | + | - | + | + | + | + | + | + | + | - | - | + | + | + | + | + | + | + | + |
| + | + | - | + | + | + | + | + | + | + | - | + | + | + | + | + | + | + | + | + |
| + | + | + | + | + | + | + | + | + | + | - | + | + | + | + | + | + | + | + | + |
| + | + | - | + | + | + | + | + | + | + | - | + | + | + | + | + | + | + | + | + |
| + | + | - | + | + | + | + | + | + | + | - | + | + | + | + | + | + | + | + | + |
| + | + | + | + | + | + | + | + | + | + | - | + | + | + | + | + | + | + | + | + |
| + | + | + | + | + | + | + | + | + | + | - | + | + | + | + | + | + | + | + | + |
| + | + | - | + | + | + | + | + | + | + | - | + | + | + | + | + | - | + | + | + |
| + | + | - | + | + | + | + | + | + | + | - | - | + | + | + | + | + | + | + | + |
| + | + | - | + | + | + | + | + | + | + | - | + | + | + | + | + | + | + | + | + |
| + | + | - | + | + | + | + | + | + | + | - | + | + | + | + | + | + | + | + | + |
| + | + | - | + | + | + | + | + | + | + | - | + | + | + | + | + | + | + | + | + |
| + | + | + | + | + | + | + | + | + | + | - | + | + | + | + | + | + | + | + | + |
| + | + | - | + | + | + | + | + | + | + | - | + | + | + | + | + | + | + | + | + |
| + | + | - | + | + | + | + | + | + | + | - | + | + | + | + | + | + | + | + | + |
| + | + | + | + | + | + | + | + | + | + | - | + | + | + | + | + | + | + | + | + |
| + | + | + | + | + | + | + | + | + | + | - | + | + | + | + | + | + | + | + | + |
| + | + | - | + | + | + | + | + | + | + | - | + | + | + | + | + | + | + | + | + |
| + | + | - | + | + | + | + | + | + | + | - | + | + | + | + | + | + | + | + | + |
| + | + | + | + | + | + | - | + | + | + | - | + | + | + | + | + | + | + | + | + |
| + | + | + | + | + | + | - | + | + | + | - | + | + | + | + | + | + | + | + | + |
| + | + | + | + | + | + | + | + | + | + | - | + | + | + | + | + | + | + | + | + |
| + | + | - | + | + | + | + | + | + | + | - | - | + | + | + | + | + | + | + | + |
| + | + | + | + | + | + | + | + | + | + | - | + | + | + | + | + | + | + | + | + |
| + | + | - | + | + | + | + | + | + | + | - | + | + | + | + | + | + | + | + | + |
| + | + | + | + | + | + | + | + | + | + | + | - | + | + | + | + | + | + | + | + |
| + | + | + | + | + | + | + | + | + | + | - | + | + | + | - | + | + | + | + | + |
| 30 | 30 | 12 | 30 | 30 | 30 | 28 | 30 | 30 | 30 | 1 | 26 | 30 | 30 | 29 | 30 | 29 | 30 | 30 | 30 |

| Cell surface components | | | | | | | | | | | | | | |
|---|---|---|---|---|---|---|---|---|---|---|---|---|---|---|
| | | | | | Heparin binding hemagglutinin | Lipoprotein | Methyltransferase | Mycolic acid transcyclopropane synthetase | MymA operon | | | PDIM (phthiocerol dimycocerosate) ar | | |
| pks | rmlA | rmlB | rmt3 | rmt4 | hbhA | lprG | mmaA4 | cmaA2 | adhD | fadD13 | sadH | Undetermined | Undetermined | Undetermined |
| + | + | + | + | - | + | + | + | + | - | + | - | + | - | - |
| + | + | + | + | - | + | + | + | + | - | + | - | + | - | - |
| + | + | + | + | - | + | + | + | + | - | + | - | + | - | - |
| + | + | + | + | - | + | + | + | + | - | + | - | + | - | - |
| + | + | + | + | - | + | + | + | + | + | + | - | + | - | - |
| + | + | + | + | - | + | + | + | + | - | + | - | + | - | - |
| + | + | + | + | - | + | + | + | + | - | + | - | + | - | - |
| + | + | + | + | - | + | + | + | + | - | + | + | + | - | - |
| + | + | + | + | - | + | + | + | + | - | + | - | + | - | - |
| + | + | + | + | - | + | + | + | + | - | + | - | + | - | - |
| + | + | + | + | - | + | + | + | + | - | + | - | + | - | - |
| + | + | + | + | - | + | + | + | + | - | + | - | + | - | - |
| + | - | + | + | - | + | + | + | + | - | + | - | + | - | - |
| + | + | + | + | - | + | + | + | + | - | + | - | + | - | - |
| + | + | + | + | - | + | + | + | + | - | + | - | + | - | - |
| + | - | + | + | - | + | + | + | + | - | + | - | + | - | - |
| + | + | + | + | - | + | + | + | + | - | + | - | + | - | - |
| + | + | + | + | - | + | + | + | + | - | + | - | + | - | - |
| + | - | + | + | - | + | + | + | + | - | + | - | + | - | - |
| + | + | + | + | - | + | + | + | + | - | + | - | + | - | - |
| + | + | + | + | - | + | + | + | + | - | + | - | + | - | - |
| + | - | + | + | - | + | + | + | + | - | + | - | + | - | - |
| + | + | + | + | - | + | + | + | + | - | + | - | + | - | - |
| + | + | + | + | - | + | + | + | + | - | + | - | + | - | - |
| + | + | + | - | + | + | + | + | + | - | + | - | - | - | - |
| + | + | + | + | - | + | + | + | + | - | + | - | + | - | - |
| + | + | + | + | - | + | + | + | + | - | + | - | + | - | - |
| + | - | + | + | - | + | + | + | + | - | + | - | + | - | - |
| + | - | + | + | + | + | + | + | + | + | + | + | + | + | + |
| + | - | + | + | - | + | + | + | + | - | + | - | + | - | - |
| 30 | 23 | 30 | 29 | 2 | 30 | 30 | 30 | 30 | 2 | 30 | 2 | 29 | 1 | 1 |

| | | | | | | | | | | | | | | | | | | | |
|---|---|---|---|---|---|---|---|---|---|---|---|---|---|---|---|---|---|---|---|
| | | | | | | | | | | | | | | | | | Copper uptake | | |
| d PGL (phenolic glycolipid) biosynthesis and transport | | | | | | | Potassium/ proton antiporter | Proximal cyclopropane synthase of alpha mycolates | Sulfolipid-1 biosynthesis and transport | | | Trehalose-recycling ABC transporter | | | | | Copper exporter | ABC transporter | |
| Undetermined | ddrA | ddrB | drrC | fadD28 | pks15/1 | pks15 | kefB | pcaA | papA2 | pks2 | stf0 | icl2 | lpqY | sugA | sugB | sugC | ctpV | irtA | irtB |
| + | + | + | + | + | - | + | + | + | - | - | + | + | + | + | + | + | + | + | + |
| + | + | + | + | + | - | + | + | + | - | - | + | + | + | + | + | + | - | + | + |
| + | + | + | + | + | - | + | + | + | - | - | + | + | + | + | + | + | - | + | + |
| + | + | + | + | + | - | + | + | + | - | - | + | + | + | + | + | + | + | + | + |
| + | + | + | + | + | - | + | + | + | - | - | + | + | + | + | + | + | + | + | + |
| + | + | + | + | + | - | + | + | + | - | - | + | + | + | + | + | + | - | + | + |
| - | + | + | + | + | - | + | + | + | - | - | + | + | + | + | + | + | - | + | + |
| - | + | + | + | + | - | + | + | + | - | - | + | + | + | + | + | + | - | + | + |
| + | + | + | + | - | - | + | + | + | - | - | + | + | + | + | + | + | - | + | + |
| - | + | + | + | + | - | + | + | + | - | - | + | + | + | + | + | + | - | + | + |
| - | + | + | + | + | - | + | + | + | - | - | + | + | + | + | + | + | + | + | + |
| - | + | + | + | + | - | + | + | + | - | - | + | + | + | + | + | + | - | + | + |
| - | + | + | + | - | - | + | + | + | - | - | + | + | + | + | + | + | - | + | + |
| - | + | + | + | + | - | + | + | + | - | - | + | + | + | + | + | + | - | + | + |
| - | + | + | + | + | - | + | + | + | - | - | + | + | + | + | + | + | - | + | + |
| - | + | + | + | - | - | + | + | + | - | - | + | + | + | + | + | + | - | + | + |
| - | + | + | + | + | - | + | + | + | - | - | + | + | + | + | + | + | - | + | + |
| - | + | + | + | + | - | + | + | + | - | - | + | + | + | + | + | + | - | + | + |
| - | + | + | + | - | - | + | + | + | - | - | + | + | + | + | + | + | - | + | + |
| - | + | + | + | - | - | + | + | + | - | - | + | + | + | + | + | + | - | + | + |
| - | + | + | + | + | - | + | + | + | - | - | + | + | + | + | + | + | - | + | + |
| - | + | + | + | - | - | + | + | + | - | - | + | + | + | + | + | + | - | + | + |
| - | + | + | + | - | - | + | + | + | - | - | + | + | + | + | + | + | - | + | + |
| - | + | + | + | - | - | + | + | + | - | - | + | + | + | + | + | + | - | + | + |
| - | + | + | + | + | - | + | + | + | - | - | + | + | + | + | + | + | - | + | + |
| - | + | + | + | + | - | + | + | + | - | - | + | + | + | + | + | + | + | + | + |
| - | + | + | + | - | - | + | + | + | - | - | + | + | + | + | + | + | - | + | + |
| - | + | + | + | - | - | + | + | + | - | - | + | + | + | + | + | + | - | + | + |
| - | + | + | + | - | + | - | + | + | + | + | + | + | + | + | + | + | + | + | + |
| - | + | + | + | - | - | + | + | + | - | - | + | + | + | + | + | + | - | + | + |
| 7 | 30 | 30 | 30 | 18 | 1 | 29 | 30 | 30 | 1 | 1 | 30 | 30 | 30 | 30 | 30 | 30 | 6 | 30 | 30 |

| Iron uptake | | | | | | | | | | | | | | | | | | |
|---|---|---|---|---|---|---|---|---|---|---|---|---|---|---|---|---|---|---|
| Heme uptake | | | Iron-dependent regulator | Mycobactin | | | | | | | | | | | | | | Heme biosynthesis (Haemophilus) |
| Undetermined | mmpL11 | mmpL3 | ideR | Undetermined | fadD33 | fadE14 | mbtA | mbtB | mbtC | mbtD | mbtE | mbtF | mbtG | mbtH | mbtI | mbtJ | mbtK | hemL |
| - | + | - | + | + | + | + | + | + | + | + | + | + | + | + | + | + | + | - |
| - | + | - | + | + | + | + | + | + | + | + | + | + | + | + | + | + | + | - |
| - | + | - | + | + | + | + | + | + | + | + | + | + | + | + | + | + | + | - |
| - | + | - | + | + | + | + | + | + | + | + | + | + | + | + | + | + | + | - |
| + | + | - | + | + | + | + | + | + | + | + | + | + | + | + | + | + | + | - |
| - | + | - | + | + | + | + | + | + | + | + | + | + | + | + | + | + | + | - |
| - | + | - | + | + | + | + | + | + | + | + | + | + | + | + | + | + | + | - |
| - | + | - | + | + | + | + | + | + | + | + | + | + | + | + | + | + | + | - |
| + | + | + | + | + | + | + | + | + | + | + | + | + | + | + | + | + | + | - |
| + | + | + | + | + | + | + | + | + | + | + | + | + | + | + | + | + | + | - |
| + | + | - | + | + | + | + | + | + | + | + | + | + | + | + | + | + | + | - |
| + | + | + | + | + | + | + | + | + | + | + | + | + | + | + | + | + | + | - |
| + | + | - | + | + | + | + | + | + | + | + | + | + | + | + | + | + | + | - |
| + | + | + | + | + | + | + | + | + | + | + | + | + | + | + | + | + | + | - |
| + | + | + | + | + | + | + | + | + | + | + | + | + | + | + | + | + | + | - |
| + | + | + | + | + | + | + | + | + | + | + | + | + | + | + | + | + | + | - |
| + | + | + | + | + | + | + | + | + | + | + | + | + | + | + | + | + | + | - |
| + | + | + | + | + | + | + | + | + | + | + | + | + | + | + | + | + | + | - |
| + | + | + | + | + | + | + | + | + | + | + | + | + | + | + | + | + | + | - |
| + | + | + | + | + | + | + | + | + | + | + | + | + | + | + | + | + | + | - |
| + | + | - | + | + | + | + | + | + | + | + | + | + | + | + | + | + | + | - |
| + | + | + | + | + | + | + | + | + | + | + | + | + | + | + | + | + | + | - |
| + | + | + | + | + | + | + | + | + | + | + | + | + | + | + | + | + | + | - |
| + | + | + | + | + | + | + | + | + | + | + | + | + | + | + | + | + | + | - |
| + | + | + | + | + | + | + | + | + | + | + | + | + | + | + | + | + | + | - |
| + | + | + | + | + | + | + | + | + | + | + | + | + | + | + | + | + | + | - |
| + | + | + | + | + | + | + | + | + | + | + | + | + | + | + | + | + | + | - |
| + | + | - | + | + | + | + | + | + | + | + | + | + | + | + | + | + | + | - |
| + | + | + | + | + | + | + | + | + | + | + | + | + | + | + | + | + | + | + |
| + | + | + | + | + | + | + | + | + | + | + | + | + | + | + | + | + | + | - |
| 23 | 30 | 18 | 30 | 30 | 30 | 30 | 30 | 30 | 30 | 30 | 30 | 30 | 30 | 30 | 30 | 30 | 30 | 1 |

| Lipid and fatty acid metabolism | | | | | | Magnesium uptake | | | | | | | | | | | | | |
|---|---|---|---|---|---|---|---|---|---|---|---|---|---|---|---|---|---|---|---|
| FAS-II | Isocitrate lyase | Lipase | Lipid phosphatase | Pantothenate synthesis | | Magnesium transport | Mce1 | | | | | | Mce2 | | | | | | |
| kasB | icl | lipF | sapM | panC | panD | mgtC | mce1A | mce1B | mce1C | mce1D | mce1E | mce1F | mce2A | mce2B | mce2C | mce2D | mce2E | mce2F | mce3A |
| + | + | - | + | + | + | + | + | + | - | + | - | + | + | + | - | + | + | + | + |
| + | + | - | + | - | + | + | + | + | - | - | - | + | + | + | + | - | + | + | + |
| + | - | + | - | + | - | + | + | - | + | - | + | + | + | + | + | + | + | + | + |
| + | + | - | + | - | + | + | + | + | - | + | - | + | + | + | - | + | + | + | + |
| + | + | - | + | - | + | + | + | + | + | + | + | + | + | + | - | + | + | + | + |
| - | + | - | + | + | + | + | + | + | - | + | - | + | + | + | + | + | + | + | + |
| + | + | - | + | + | + | + | + | + | - | - | - | + | + | + | - | - | - | + | + |
| + | + | - | + | - | + | + | + | + | + | + | + | + | + | + | - | + | + | + | + |
| + | + | - | + | + | + | + | + | + | + | + | - | + | + | + | + | + | + | + | + |
| + | + | - | + | + | + | + | + | + | + | + | - | + | + | + | + | + | + | + | + |
| - | + | - | + | + | + | + | + | + | + | + | - | + | + | + | + | + | + | + | + |
| + | + | - | + | + | + | + | + | + | + | + | - | + | + | + | + | + | + | + | + |
| + | + | - | + | + | + | + | + | + | + | + | - | + | + | + | + | + | + | + | + |
| + | + | - | + | + | + | + | + | + | + | + | - | + | + | + | + | + | + | + | + |
| + | + | - | + | + | + | + | + | + | + | + | - | + | + | + | + | + | + | + | + |
| + | + | - | + | + | + | + | + | + | + | + | - | + | + | + | + | + | + | + | + |
| + | + | - | + | + | + | + | + | + | + | + | - | + | + | + | + | + | + | + | + |
| + | + | - | + | + | + | + | + | + | + | + | - | + | + | + | + | + | + | + | + |
| + | + | - | + | + | + | + | + | + | + | + | - | + | + | + | + | + | + | + | + |
| + | + | - | + | + | + | + | + | + | + | + | - | + | + | + | + | + | + | + | + |
| + | + | - | + | + | + | + | + | + | + | + | - | + | + | + | + | + | + | + | + |
| + | + | - | + | + | + | + | + | + | + | + | - | + | + | + | + | + | + | + | + |
| + | + | - | + | + | + | + | + | + | + | + | - | + | + | + | + | + | + | + | + |
| + | + | - | + | + | + | + | + | + | + | + | - | + | + | + | + | + | + | + | + |
| + | + | - | + | + | + | + | + | + | + | + | - | + | + | + | + | + | + | + | + |
| + | + | - | + | + | + | + | + | + | + | + | - | + | + | + | + | + | + | + | + |
| + | + | - | + | + | + | + | + | + | + | + | - | + | + | + | + | + | + | + | + |
| + | + | - | + | + | + | + | + | + | + | + | - | + | + | + | + | + | + | + | + |
| + | + | + | + | + | + | + | + | + | + | + | + | + | + | + | + | + | + | + | + |
| + | + | - | + | + | + | + | + | + | + | + | + | + | + | + | + | + | + | + | + |
| 28 | 29 | 2 | 29 | 26 | 29 | 30 | 30 | 29 | 25 | 27 | 5 | 30 | 30 | 30 | 25 | 28 | 29 | 30 | 30 |

| Mammalian cell entry (mce) operons | | | | | | | | | | | | | | | | | | | |
|---|---|---|---|---|---|---|---|---|---|---|---|---|---|---|---|---|---|---|---|
| Mce3 | | | | | Mce4 | | | | | | Mce5 | | | | | | Mc | | |
| mce3B | mce3C | mce3D | mce3E | mce3F | mce4A | mce4B | mce4C | mce4D | mce4E | mce4F | mce5A | mce5B | mce5C | mce5D | mce5E | mce5F | mce7A | mce7B | mce7C |
| + | + | + | + | + | + | + | + | - | + | - | + | + | + | + | + | + | + | + | + |
| + | + | + | + | + | + | + | + | - | + | - | + | + | + | + | + | + | + | + | + |
| + | + | + | + | + | + | + | + | + | + | + | + | + | + | + | + | - | + | + | + |
| + | + | + | + | + | + | + | + | - | + | + | + | + | + | + | + | + | + | + | + |
| + | + | + | - | + | + | + | + | - | + | + | + | + | + | - | + | + | + | + | + |
| + | + | + | + | + | + | + | + | - | + | + | + | + | + | + | + | + | + | + | + |
| + | + | + | + | + | + | + | + | + | + | + | + | + | + | + | + | + | + | + | + |
| + | + | + | - | + | + | + | + | + | + | + | + | + | + | + | + | + | + | + | + |
| + | + | + | + | + | + | + | + | + | + | + | + | + | + | + | + | + | + | + | + |
| + | + | + | + | + | + | + | + | + | + | + | + | + | + | + | + | - | + | + | + |
| + | + | + | + | + | + | + | + | + | + | + | + | + | + | + | + | + | + | + | + |
| + | + | + | + | + | + | + | + | + | + | + | + | + | + | + | + | - | + | + | + |
| + | + | + | + | + | + | + | + | + | + | + | + | + | + | + | + | + | + | + | + |
| + | + | + | + | + | + | + | + | + | + | + | + | + | + | + | + | - | + | + | + |
| + | + | + | + | + | + | + | - | + | + | + | + | + | + | + | + | + | + | + | + |
| + | + | + | + | + | + | + | + | + | + | + | + | + | + | + | + | + | + | + | + |
| + | + | + | + | + | + | + | + | + | + | + | + | + | + | + | + | - | + | + | + |
| + | + | + | + | + | + | + | + | + | + | + | + | + | + | + | + | - | + | + | + |
| + | + | + | + | + | + | + | + | + | + | + | + | + | + | + | + | + | + | + | + |
| + | + | + | + | + | + | + | + | + | + | + | + | + | + | + | + | + | + | + | + |
| + | + | + | + | + | + | + | + | + | + | + | + | + | + | + | + | - | + | + | + |
| + | + | + | + | + | + | + | + | + | + | + | + | + | + | + | + | + | + | + | + |
| + | + | + | + | + | + | + | + | + | + | + | + | + | + | + | + | + | + | + | + |
| + | + | + | + | + | + | + | + | + | + | + | + | + | + | + | + | + | + | + | + |
| + | + | + | + | + | + | + | + | + | + | + | + | + | + | + | + | + | + | + | + |
| + | + | + | + | + | + | + | + | + | + | + | + | + | + | + | + | + | + | + | + |
| + | + | + | + | + | + | + | + | + | + | + | + | + | + | + | + | + | + | + | + |
| + | + | + | + | + | + | + | + | + | + | + | + | + | + | + | + | + | + | + | + |
| + | + | + | + | + | + | + | + | + | + | + | + | + | + | + | + | + | + | + | + |
| + | + | + | + | + | + | + | + | + | + | + | + | + | + | + | + | + | + | + | + |
| 30 | 30 | 30 | 28 | 30 | 30 | 30 | 29 | 25 | 30 | 28 | 30 | 30 | 30 | 29 | 30 | 23 | 30 | 30 | 30 |

| | | | | | | | | | Manganese uptake | Phagosome arresting | | | Protease | | | |
|---|---|---|---|---|---|---|---|---|---|---|---|---|---|---|---|---|
| e7 | | | Mce9 | | | | | | P(1)-type Mn2+ transporting ATPase | Nucleoside diphosphate kinase | PE family protein | Tyrosine phosphatase | Proteasome associated proteins | | Zn2+ metallophrotease | (p)ppGpp synthesis and hydrolysis |
| mce7D | mce7E | mce7F | mce9A | mce9B | mce9C | mce9D | mce9E | mce9F | ctpC | ndk | PE_PGRS30 | ptpA | mpa | pafA | zmp1 | relA |
| + | + | + | + | + | + | + | + | + | + | + | - | + | + | + | + | + |
| + | + | + | - | + | + | + | + | + | + | + | - | + | + | + | + | + |
| + | + | - | + | + | + | + | + | + | + | - | + | + | + | + | + | + |
| + | + | + | - | + | + | + | + | + | + | + | - | + | + | + | + | + |
| + | + | + | + | + | + | + | + | + | + | + | - | + | + | + | + | + |
| + | + | + | + | + | + | + | + | + | + | + | - | + | + | + | + | + |
| + | + | + | + | + | + | + | + | + | + | + | - | + | + | + | + | + |
| + | + | + | + | + | + | + | + | + | + | + | - | + | + | + | + | + |
| + | + | + | + | + | + | + | + | + | + | + | - | + | + | + | + | + |
| + | + | + | + | + | + | + | + | + | + | + | - | + | + | + | + | + |
| + | + | + | + | + | + | + | + | + | + | + | - | + | + | + | + | + |
| + | + | + | + | + | + | + | + | + | + | + | - | + | + | + | + | + |
| + | + | + | + | + | + | + | + | + | + | + | - | + | + | + | + | + |
| + | + | + | + | + | + | + | + | + | + | + | - | + | + | + | + | + |
| + | + | + | + | + | + | + | + | + | + | + | - | + | + | + | + | + |
| + | + | + | + | + | + | + | + | + | + | + | - | + | + | + | + | + |
| + | + | + | + | + | + | + | + | + | + | + | - | + | + | + | + | + |
| + | + | + | + | + | + | + | + | + | + | + | - | + | + | + | + | + |
| + | + | + | + | + | + | + | + | + | + | + | - | + | + | + | + | + |
| + | + | + | + | + | + | + | + | + | + | + | - | + | + | + | + | + |
| + | + | + | + | + | + | + | + | + | + | + | - | + | + | + | + | + |
| + | + | + | + | + | + | + | + | + | + | + | - | + | + | + | + | + |
| + | + | + | + | + | + | + | + | + | + | + | - | + | + | + | + | + |
| + | + | + | + | + | + | + | + | + | + | + | - | + | + | + | + | + |
| + | + | + | + | + | + | + | + | + | + | + | - | + | + | + | + | + |
| + | + | + | + | + | + | + | + | + | + | + | - | + | + | + | + | + |
| + | + | + | + | + | + | + | + | + | + | + | - | + | + | + | + | + |
| + | + | + | + | + | + | + | + | + | + | + | - | + | + | + | + | + |
| + | + | + | + | + | + | + | + | + | + | + | - | + | + | + | + | + |
| + | + | + | + | + | + | + | + | + | + | + | - | + | + | + | + | + |
| 30 | 30 | 29 | 28 | 30 | 30 | 30 | 30 | 30 | 30 | 29 | 1 | 30 | 30 | 30 | 30 | 30 |

| Regulation | | | | | | | | | | | | | | | | | | | |
|---|---|---|---|---|---|---|---|---|---|---|---|---|---|---|---|---|---|---|---|
| DevR/S | | MosR | MprA/B | | PhoP/R | | PrrA/B | | RegX3 | SenX3 | Sigma A | Sigma D | Sigma E | Sigma F | Sigma H | Sigma L | Sigma M | WhiB3 | 19-kD protein |
| devR/dosR | devS | mosR | mprA | mprB | phoP | phoR | prrA | prrB | regX3 | senX3 | sigA/rpoV | sigD | sigE | sigF | sigH | sigL | sigM | whiB3 | lpqH |
| + | + | - | + | + | + | + | + | + | + | + | + | + | + | + | + | + | - | + | - |
| + | + | - | + | + | + | + | + | + | + | + | + | - | + | + | + | + | + | + | + |
| + | - | + | + | + | + | + | + | + | + | + | + | + | + | + | + | + | + | + | - |
| + | + | - | + | + | + | + | + | + | + | + | + | + | - | + | + | + | + | + | - |
| + | + | - | + | + | + | + | + | + | + | + | + | + | + | + | + | + | + | + | - |
| + | + | - | + | + | + | + | + | + | + | + | + | + | + | + | + | + | + | + | + |
| + | + | - | + | + | + | + | + | + | + | + | + | + | + | + | + | + | + | + | - |
| + | + | - | + | + | + | + | + | + | + | + | + | + | + | + | + | + | + | + | - |
| + | + | - | + | + | + | + | + | + | + | + | + | + | + | + | + | + | + | + | - |
| + | + | - | + | + | + | + | + | + | + | + | + | + | + | + | + | + | + | + | + |
| + | + | - | + | + | + | + | + | + | + | + | + | + | + | + | + | + | + | + | - |
| + | + | - | + | + | + | + | + | + | + | + | + | + | + | + | + | + | - | + | - |
| + | + | - | + | + | + | + | + | + | + | + | + | + | + | + | + | + | + | + | - |
| + | + | - | + | + | + | + | + | + | + | + | + | + | + | + | + | + | + | + | + |
| + | + | - | + | + | + | + | + | + | + | + | + | + | + | + | + | + | + | + | - |
| + | + | - | + | + | + | + | + | + | + | + | + | + | + | + | + | + | + | + | - |
| + | + | - | + | + | + | + | + | + | + | + | + | + | + | + | + | + | + | + | + |
| + | + | - | + | + | + | + | + | + | + | + | + | + | + | + | + | + | - | + | - |
| + | + | - | + | + | + | + | + | + | + | + | + | + | + | + | + | + | + | + | - |
| + | + | - | + | + | + | + | + | + | + | + | + | + | + | + | + | + | + | + | + |
| + | + | - | + | + | + | + | + | + | + | + | + | + | + | + | + | + | + | + | - |
| + | + | - | + | + | + | + | + | + | + | + | + | + | + | + | + | + | + | + | - |
| + | + | - | + | + | + | + | + | + | + | + | + | + | + | + | + | + | + | + | + |
| + | + | - | + | + | + | + | + | + | + | + | + | + | + | + | + | + | + | + | + |
| + | + | - | + | + | + | + | + | + | + | + | + | + | + | + | + | + | + | + | + |
| + | + | - | + | + | + | + | + | + | + | + | + | + | + | + | + | + | + | + | + |
| + | + | - | + | + | + | + | + | + | + | + | + | + | + | + | + | + | + | + | + |
| + | + | - | + | + | + | + | + | + | + | + | + | + | + | + | + | + | + | + | - |
| + | + | - | + | + | + | + | + | + | + | + | + | + | + | + | + | + | + | + | - |
| + | + | - | + | + | + | + | + | + | + | + | + | + | + | + | + | + | + | + | + |
| 30 | 29 | 1 | 30 | 30 | 30 | 30 | 30 | 30 | 30 | 30 | 30 | 29 | 29 | 30 | 30 | 30 | 27 | 30 | 12 |

| Secreted proteins | | | | | | | | | | | | | | | | | | | | |
|---|---|---|---|---|---|---|---|---|---|---|---|---|---|---|---|---|---|---|---|---|
| Alpha-crystallin | Antigen 85 complex | | | Enhanced intracellular survival protein | Protein kinase G | Accessory secretion factor | | | | | | ESX-2 (T7SS) | | | | | | | | |
| hspX | fbpA | fbpB | fbpC | eis | pknG | secA2 | eccA1 | espI | espL | espR | mycP1 | Undetermined | PE36 | PPE69 | eccA2 | eccB2 | eccC2 | eccD2 | eccE2 | espG2 |
| - | + | + | + | + | + | + | - | - | - | + | - | + | + | + | + | + | + | + | + | + |
| - | + | + | + | + | + | + | - | - | - | + | - | + | + | + | + | + | + | + | + | + |
| + | + | + | + | + | + | - | - | - | + | - | + | + | + | + | + | + | + | + | + | + |
| - | + | + | + | + | + | + | - | - | - | + | - | + | + | + | + | + | + | + | + | + |
| - | + | + | + | + | + | + | + | - | - | + | - | + | + | + | + | + | + | + | + | + |
| - | + | + | + | + | + | + | - | - | - | + | - | + | + | + | + | + | + | + | + | + |
| - | + | + | + | + | + | + | - | - | - | + | - | + | + | + | + | + | + | + | + | + |
| - | + | + | + | + | + | + | - | - | - | + | - | + | + | + | + | + | + | + | + | + |
| - | + | + | + | + | + | + | - | - | - | + | - | + | + | + | + | + | + | + | + | + |
| - | + | + | + | + | + | + | - | - | - | + | - | + | + | + | + | + | + | + | + | + |
| - | + | + | + | + | + | + | - | - | - | + | - | + | + | + | + | + | + | + | + | + |
| - | + | + | + | + | + | + | - | - | - | - | - | + | + | + | + | + | + | + | + | + |
| - | + | + | + | + | + | + | - | - | - | + | - | + | + | + | + | + | + | + | + | + |
| - | + | + | + | + | + | + | - | - | - | + | - | + | + | + | + | + | + | + | + | + |
| - | + | + | + | + | + | + | - | - | - | + | - | + | + | + | + | + | + | + | + | + |
| - | + | + | + | + | + | + | - | - | - | + | - | + | + | + | + | + | + | + | + | + |
| - | + | + | + | + | + | + | - | - | - | + | - | + | + | + | + | + | + | + | + | + |
| - | + | + | + | + | + | + | - | - | - | + | - | + | + | + | + | + | + | + | + | + |
| - | + | + | + | + | + | + | - | - | - | + | - | + | + | + | + | + | + | + | + | + |
| - | + | + | + | + | + | + | - | - | - | + | - | + | + | + | + | + | + | + | + | + |
| - | + | + | + | + | + | + | + | - | - | + | - | + | + | + | + | + | + | + | + | + |
| - | + | + | + | + | + | + | - | - | - | + | - | + | + | + | + | + | + | + | + | + |
| - | + | + | + | + | + | + | - | - | - | + | - | + | + | + | + | + | - | + | + | + |
| - | + | + | + | + | + | + | - | - | - | + | - | + | + | + | + | + | - | + | + | + |
| - | + | + | - | + | + | + | - | - | - | + | - | + | + | + | + | + | + | + | + | + |
| - | + | + | + | + | + | + | - | - | - | + | - | + | + | + | + | + | + | + | + | + |
| - | + | + | + | + | + | + | - | - | - | + | - | + | + | + | + | + | + | + | + | + |
| - | + | + | + | + | + | + | - | - | - | + | - | + | + | + | + | + | + | + | + | + |
| + | + | + | + | + | + | + | - | + | - | + | - | + | + | + | + | + | + | + | + | + |
| - | + | + | + | + | + | + | - | - | - | + | - | + | + | + | + | + | + | + | + | + |
| 2 | 30 | 30 | 29 | 30 | 30 | 29 | 2 | 1 | 1 | 28 | 1 | 30 | 30 | 30 | 30 | 30 | 28 | 30 | 30 | 30 |

| Secretion system | | | | | | | | | | | | | | | | | | | | |
|---|---|---|---|---|---|---|---|---|---|---|---|---|---|---|---|---|---|---|---|---|
| | | | ESX-3 (T7SS) | | | | | | | | | | | ESX-4 (T7SS) | | | | | | |
| esxC | esxD | mycP2 | PE5 | PPE4 | eccA3 | eccB3 | eccC3 | eccD3 | eccE3 | espG3 | esxG | esxH | mycP3 | Undetermined | eccB4 | eccC4 | eccD4 | esxT | esxU | mycP4 |
| + | + | - | - | + | + | + | + | + | + | + | + | + | - | - | + | + | - | + | + | - |
| + | + | - | - | + | + | + | + | + | + | + | + | + | - | + | - | + | - | + | + | - |
| + | - | - | + | + | + | + | + | + | + | + | + | - | - | - | + | - | + | + | - | + |
| + | + | - | + | + | + | + | + | + | + | + | + | + | - | - | - | + | - | + | + | - |
| + | + | - | - | + | + | + | + | + | + | + | + | + | + | - | - | + | - | + | + | - |
| + | + | - | + | + | + | + | + | + | + | + | + | + | + | - | - | + | - | + | + | - |
| + | + | - | - | + | + | + | + | + | + | + | + | + | - | + | - | + | - | + | + | - |
| + | + | + | - | + | + | + | + | + | + | + | + | + | + | - | - | + | - | + | + | - |
| + | + | + | + | + | + | + | + | + | + | + | + | + | + | + | + | + | + | + | + | + |
| + | + | + | + | + | + | + | + | + | + | + | + | + | + | + | + | + | + | + | + | + |
| + | + | + | + | + | + | + | + | + | + | + | + | + | + | + | + | + | + | + | + | + |
| + | + | + | + | + | + | + | + | + | + | + | + | + | + | + | + | + | + | + | + | + |
| + | + | + | + | + | + | + | + | + | + | + | + | + | + | + | + | + | + | + | + | + |
| + | + | + | + | + | + | + | + | + | + | + | + | + | + | + | + | + | + | + | + | + |
| + | + | + | + | + | + | + | + | + | + | + | + | + | + | + | + | + | + | - | + | + |
| + | + | + | + | + | + | + | + | + | + | + | + | + | + | + | + | + | + | + | + | + |
| + | + | + | + | + | + | + | + | + | + | + | + | + | + | + | + | + | + | + | + | + |
| + | + | + | + | + | + | + | + | + | + | + | + | + | + | + | + | + | + | + | + | + |
| + | + | + | + | + | + | + | + | + | + | + | + | + | + | + | + | + | + | + | + | + |
| + | + | + | + | + | + | + | + | + | + | + | + | + | + | + | + | + | + | + | + | + |
| + | + | + | + | + | + | + | + | + | + | + | + | + | + | + | + | + | + | + | + | + |
| + | + | + | + | + | + | + | + | + | + | + | + | + | + | + | + | + | + | + | + | + |
| + | + | + | + | + | + | + | + | + | + | + | + | + | + | + | + | + | + | + | + | + |
| + | + | + | + | + | + | + | + | + | + | + | + | + | + | + | + | + | + | + | + | + |
| + | + | + | + | + | + | + | + | + | + | + | + | + | + | + | + | + | + | + | + | + |
| + | + | + | + | + | + | + | + | + | + | + | + | + | + | + | + | + | + | + | + | + |
| + | + | + | + | + | + | + | + | + | + | + | + | + | + | + | + | + | + | + | + | + |
| + | + | + | + | + | + | + | + | + | + | + | + | + | + | + | + | + | + | + | + | + |
| + | + | + | + | + | + | + | + | + | + | + | + | + | + | + | + | + | + | + | + | + |
| + | + | + | + | + | + | + | + | + | + | + | + | + | + | + | + | + | + | + | + | + |
| 30 | 29 | 23 | 25 | 30 | 30 | 30 | 30 | 30 | 30 | 30 | 30 | 29 | 25 | 24 | 24 | 29 | 23 | 29 | 29 | 23 |

| ESX-5 (T7SS) | | | | | | | | | | | | | | | | | T6SS-II (Klebsiella) | AhpC |
|---|---|---|---|---|---|---|---|---|---|---|---|---|---|---|---|---|---|---|
| Undetermined | Undetermined | PE18 | PE19 | PPE25 | PPE26 | PPE41 | cyp143 | eccA5 | eccB5 | eccCa5 | eccCb5 | eccD5 | eccE5 | esxM | esxN | mycP5 | - | ahpC |
| + | + | - | + | + | + | - | + | + | + | - | + | + | + | + | + | + | - | + |
| + | + | + | + | - | - | - | + | + | + | - | + | + | + | + | + | + | - | + |
| + | + | + | + | + | - | + | + | + | - | + | + | + | + | + | + | + | - | + |
| + | + | + | + | + | - | - | + | + | + | - | + | + | + | + | + | + | - | + |
| + | + | + | + | + | + | - | + | + | + | - | + | + | + | + | + | + | - | + |
| + | + | + | + | + | + | - | + | + | + | - | + | + | + | + | + | + | - | + |
| + | + | - | - | + | + | - | + | + | + | - | + | + | + | + | + | + | - | + |
| + | + | + | - | + | + | - | + | + | + | - | + | + | + | + | + | + | - | + |
| + | + | + | - | - | + | - | + | + | + | - | + | + | + | + | + | + | - | + |
| + | + | + | + | - | + | - | + | + | + | - | + | + | + | + | + | + | - | + |
| + | + | + | - | + | - | - | + | + | + | - | + | + | + | + | + | + | - | + |
| + | + | + | + | - | + | - | + | + | + | - | + | + | + | + | + | + | - | + |
| + | + | + | - | - | - | - | - | + | - | - | - | + | + | + | + | + | - | + |
| + | + | + | + | + | + | - | + | + | + | - | + | + | + | + | + | + | - | + |
| + | + | + | + | - | + | - | + | + | + | - | + | + | + | + | + | + | - | + |
| + | + | + | - | - | + | - | + | + | + | - | + | + | + | + | + | + | - | + |
| + | + | + | + | - | + | - | + | + | + | - | + | + | + | + | + | + | - | + |
| + | + | + | + | - | + | - | + | + | + | - | + | + | + | + | + | + | - | + |
| + | + | + | - | - | + | - | + | + | + | - | + | + | + | + | + | + | - | + |
| + | + | + | + | - | + | - | + | + | + | - | + | + | + | + | + | + | - | + |
| + | + | + | - | - | + | - | + | + | + | - | + | + | + | + | + | + | - | + |
| + | + | + | - | - | - | - | + | + | + | - | + | + | + | + | + | + | - | + |
| + | + | + | - | - | - | - | + | + | + | - | + | + | + | + | + | + | - | + |
| + | + | + | - | - | - | - | + | + | + | - | + | + | + | + | + | + | - | + |
| + | + | + | - | - | + | - | + | + | + | - | + | + | + | + | + | + | - | + |
| + | + | + | - | + | + | - | + | + | + | - | + | + | + | + | + | + | - | + |
| + | + | + | + | - | + | - | + | + | + | - | + | + | + | + | + | + | - | + |
| + | + | + | - | - | - | - | + | + | + | - | + | + | + | + | + | + | - | + |
| + | + | + | - | - | - | - | + | + | + | - | + | + | + | + | + | + | + | + |
| + | + | + | - | - | + | - | + | + | + | - | + | + | + | + | + | + | + | + |
| 30 | 30 | 28 | 14 | 10 | 20 | 1 | 29 | 30 | 28 | 1 | 29 | 30 | 30 | 30 | 30 | 30 | 2 | 30 |

| Stress adaptation | | | Toxin | Immune evasion | Adherence | | Antiphagocytosis | Enzyme | Immu | |
|---|---|---|---|---|---|---|---|---|---|---|
| Catalase-peroxidase | Cu | Iron-cofactored SOD | Mycolactone | Capsule (Acinetobacter) | GroEL (Clostridium) | Streptococcal plasmin receptor/GAPDH (Streptococcus) | Capsule (Klebsiella) | Streptococcal enolase (Streptococcus) | Capsule (Acinetobacter) | Capsule (Streptococcus) |
| katG | sodC | sodA | mlsB | - | groEL | plr/gapA | gnd | eno | - | rmlB |
| + | + | + | - | + | - | - | - | - | - | - |
| + | + | + | - | - | - | - | - | - | - | - |
| + | + | - | - | - | - | - | - | - | - | - |
| + | + | + | - | - | - | - | - | - | - | - |
| + | + | + | - | - | - | - | - | - | - | - |
| + | + | + | - | - | - | - | - | - | - | - |
| + | + | + | - | - | - | - | - | - | - | - |
| + | + | + | - | - | - | - | - | - | - | - |
| + | + | + | - | - | - | - | - | - | - | - |
| + | + | + | - | - | - | - | - | - | - | - |
| + | + | + | - | - | - | - | - | - | - | - |
| + | + | - | - | - | - | - | - | - | - | - |
| + | + | + | - | + | + | + | + | + | + | + |
| + | + | + | - | - | - | - | - | - | - | - |
| + | + | + | - | - | - | - | - | - | - | - |
| + | + | + | - | - | - | - | - | - | - | - |
| + | + | + | - | - | - | - | - | - | - | - |
| + | + | + | - | - | - | - | - | - | - | - |
| + | + | + | - | - | - | - | - | - | - | - |
| + | + | + | - | - | - | - | - | - | - | - |
| + | + | + | - | - | - | - | - | - | - | - |
| + | + | + | - | - | - | - | - | - | - | - |
| + | + | + | - | - | - | - | - | - | - | - |
| + | + | + | - | - | - | - | - | - | - | - |
| + | + | + | - | - | - | - | - | - | - | - |
| + | + | + | - | - | - | - | - | - | - | - |
| + | + | + | - | - | - | - | - | - | - | - |
| + | + | + | - | - | - | - | - | - | - | - |
| + | + | + | - | - | + | + | + | + | + | + |
| + | + | - | + | - | - | - | - | - | - | - |
| 30 | 30 | 27 | 1 | 2 | 2 | 2 | 2 | 2 | 2 | 2 |

| ne evasion | | Nutritional virulence | Other adhesion-related proteins | Total number of expressed virulrnce factor |
|---|---|---|---|---|
| Exopolysaccharide (Haemophilus) | LOS (Campylobacter) | Pyrimidine biosynthesis (Francisella) | EF-Tu (Mycoplasma) | |
| pgi | gmhA | carB | tuf | |
| - | - | - | - | 190 |
| - | - | - | - | 186 |
| - | - | - | - | 188 |
| - | - | - | - | 187 |
| - | - | - | - | 194 |
| - | - | - | - | 193 |
| - | - | - | - | 185 |
| - | - | - | - | 192 |
| - | - | - | - | 200 |
| - | - | - | - | 201 |
| - | - | - | - | 197 |
| - | - | - | - | 195 |
| + | + | + | + | 203 |
| - | - | - | - | 201 |
| - | - | - | - | 198 |
| - | - | - | - | 198 |
| - | - | - | - | 200 |
| - | - | - | - | 198 |
| - | - | - | - | 198 |
| - | - | - | - | 201 |
| - | - | - | - | 198 |
| - | - | - | - | 196 |
| - | - | - | - | 197 |
| - | - | - | - | 197 |
| - | - | - | - | 199 |
| - | - | - | - | 201 |
| - | - | - | - | 201 |
| - | - | - | - | 195 |
| + | + | + | + | 222 |
| - | - | - | - | 200 |
| 2 | 2 | 2 | 2 | 0 |
