## Supplementary Tables for "Genomic features of *Mycobacterium avium* subsp. *hominissuis* isolated from pigs in Japan": Supplementary_Table 5_1st response.pdf

| | RGI criteria | ARO term | SNP | Detection criteria | AMR gene family | Drug class | Resistance Mechanism | Identify of Matching Region (%) | Length of Reference Sequence (%) |
|---|---|---|---|---|---|---|---|---|---|
| GM5 | Strict | mtrA | | protein homolog model | resistance-nodulation-cell division (RND) antibiotic efflux pump | macrolide antibiotic, penam | antibiotic efflux | 99.56 | 100 |
| | Strict | RbpA | | protein homolog model | RbpA bacterial RNA polymerase-binding protein | rifamycin antibiotic | antibiotic target protection | 90.99 | 97.37 |
| | Strict | Mycobacterium tuberculosis intrinsic murA conferring resistance to fosfomycin | C117D | protein variant model | murA transferase | fosfomycin | antibiotic target alteration | 97.32 | 98.56 |
| GM10 | Strict | mtrA | | protein homolog model | resistance-nodulation-cell division (RND) antibiotic efflux pump | macrolide antibiotic, penam | antibiotic efflux | 99.56 | 100 |
| | Strict | RbpA | | protein homolog model | RbpA bacterial RNA polymerase-binding protein | rifamycin antibiotic | antibiotic target protection | 90.99 | 97.37 |
| | Strict | Mycobacterium tuberculosis intrinsic murA conferring resistance to fosfomycin | C117D | protein variant model | murA transferase | fosfomycin | antibiotic target alteration | 97.32 | 98.56 |
| GM12 | Strict | mtrA | | protein homolog model | resistance-nodulation-cell division (RND) antibiotic efflux pump | macrolide antibiotic, penam | antibiotic efflux | 99.56 | 100 |
| | Strict | RbpA | | protein homolog model | RbpA bacterial RNA polymerase-binding protein | rifamycin antibiotic | antibiotic target protection | 90.99 | 97.37 |
| | Strict | Mycobacterium tuberculosis intrinsic murA conferring resistance to fosfomycin | C117D | protein variant model | murA transferase | fosfomycin | antibiotic target alteration | 97.32 | 98.56 |
| GM16 | Strict | mtrA | | protein homolog model | resistance-nodulation-cell division (RND) antibiotic efflux pump | macrolide antibiotic, penam | antibiotic efflux | 99.56 | 100 |
| | Strict | RbpA | | protein homolog model | RbpA bacterial RNA polymerase-binding protein | rifamycin antibiotic | antibiotic target protection | 90.99 | 97.37 |
| | Strict | Mycobacterium tuberculosis intrinsic murA conferring resistance to fosfomycin | C117D | protein variant model | murA transferase | fosfomycin | antibiotic target alteration | 97.08 | 98.56 |
| GM17 | Strict | mtrA | | protein homolog model | resistance-nodulation-cell division (RND) antibiotic efflux pump | macrolide antibiotic, penam | antibiotic efflux | 99.56 | 100 |
| | Strict | RbpA | | protein homolog model | RbpA bacterial RNA polymerase-binding protein | rifamycin antibiotic | antibiotic target protection | 90.99 | 97.37 |
| | Strict | Mycobacterium tuberculosis intrinsic murA conferring resistance to fosfomycin | C117D | protein variant model | murA transferase | fosfomycin | antibiotic target alteration | 97.32 | 98.56 |
| GM21 | Strict | mtrA | | protein homolog model | resistance-nodulation-cell division (RND) antibiotic efflux pump | macrolide antibiotic, penam | antibiotic efflux | 99.56 | 100 |
| | Strict | RbpA | | protein homolog model | RbpA bacterial RNA polymerase-binding protein | rifamycin antibiotic | antibiotic target protection | 90.99 | 97.37 |
| | Strict | Mycobacterium tuberculosis intrinsic murA conferring resistance to fosfomycin | C117D | protein variant model | murA transferase | fosfomycin | antibiotic target alteration | 97.32 | 98.56 |
| GM32 | Strict | mtrA | | protein homolog model | resistance-nodulation-cell division (RND) antibiotic efflux pump | macrolide antibiotic, penam | antibiotic efflux | 99.56 | 100 |
| | Strict | RbpA | | protein homolog model | RbpA bacterial RNA polymerase-binding protein | rifamycin antibiotic | antibiotic target protection | 90.99 | 97.37 |
| | Strict | Mycobacterium tuberculosis intrinsic murA conferring resistance to fosfomycin | C117D | protein variant model | murA transferase | fosfomycin | antibiotic target alteration | 97.32 | 98.56 |
| GM44 | Strict | mtrA | | protein homolog model | resistance-nodulation-cell division (RND) antibiotic efflux pump | macrolide antibiotic, penam | antibiotic efflux | 99.56 | 100 |
| | Strict | RbpA | | protein homolog model | RbpA bacterial RNA polymerase-binding protein | rifamycin antibiotic | antibiotic target protection | 90.99 | 97.37 |
| | Strict | Mycobacterium tuberculosis intrinsic murA conferring resistance to fosfomycin | C117D | protein variant model | murA transferase | fosfomycin | antibiotic target alteration | 97.32 | 98.56 |
| OCU467 | Strict | mtrA | | protein homolog model | resistance-nodulation-cell division (RND) antibiotic efflux pump | macrolide antibiotic, penam | antibiotic efflux | 99.56 | 100 |
| | Strict | RbpA | | protein homolog model | RbpA bacterial RNA polymerase-binding protein | rifamycin antibiotic | antibiotic target protection | 90.99 | 97.37 |
| | Strict | Mycobacterium tuberculosis intrinsic murA conferring resistance to fosfomycin | C117D | protein variant model | murA transferase | fosfomycin | antibiotic target alteration | 97.32 | 98.56 |
| OCU468 | Strict | RbpA | | protein homolog model | RbpA bacterial RNA polymerase-binding protein | rifamycin antibiotic | antibiotic target protection | 90.99 | 97.37 |
| | Strict | Mycobacterium tuberculosis intrinsic murA conferring resistance to fosfomycin | C117D | protein variant model | murA transferase | fosfomycin | antibiotic target alteration | 97.32 | 98.56 |
| | Strict | mtrA | | protein homolog model | resistance-nodulation-cell division (RND) antibiotic efflux pump | macrolide antibiotic, penam | antibiotic efflux | 99.12 | 100 |
| OCU469 | Strict | RbpA | | protein homolog model | RbpA bacterial RNA polymerase-binding protein | rifamycin antibiotic | antibiotic target protection | 90.99 | 97.37 |
| | Strict | Mycobacterium tuberculosis intrinsic murA conferring resistance to fosfomycin | C117D | protein variant model | murA transferase | fosfomycin | antibiotic target alteration | 97.32 | 98.56 |
| | Strict | mtrA | | protein homolog model | resistance-nodulation-cell division (RND) antibiotic efflux pump | macrolide antibiotic, penam | antibiotic efflux | 99.12 | 100 |
| OCU470 | Strict | mtrA | | protein homolog model | resistance-nodulation-cell division (RND) antibiotic efflux pump | macrolide antibiotic, penam | antibiotic efflux | 99.56 | 100 |
| | Strict | RbpA | | protein homolog model | RbpA bacterial RNA polymerase-binding protein | rifamycin antibiotic | antibiotic target protection | 90.99 | 97.37 |
| | Strict | Mycobacterium tuberculosis intrinsic murA conferring resistance to fosfomycin | C117D | protein variant model | murA transferase | fosfomycin | antibiotic target alteration | 97.32 | 98.56 |
| OCU471 | Strict | mtrA | | protein homolog model | resistance-nodulation-cell division (RND) antibiotic efflux pump | macrolide antibiotic, penam | antibiotic efflux | 99.56 | 100 |
| | Strict | RbpA | | protein homolog model | RbpA bacterial RNA polymerase-binding protein | rifamycin antibiotic | antibiotic target protection | 90.99 | 97.37 |
| | Strict | Mycobacterium tuberculosis intrinsic murA conferring resistance to fosfomycin | C117D | protein variant model | murA transferase | fosfomycin | antibiotic target alteration | 97.32 | 98.56 |
| OCU472 | Strict | Mycobacterium tuberculosis intrinsic murA conferring resistance to fosfomycin | C117D | protein variant model | murA transferase | fosfomycin | antibiotic target alteration | 97.32 | 98.56 |
| | Strict | RbpA | | protein homolog model | RbpA bacterial RNA polymerase-binding protein | rifamycin antibiotic | antibiotic target protection | 90.99 | 97.37 |
| | Strict | mtrA | | protein homolog model | resistance-nodulation-cell division (RND) antibiotic efflux pump | macrolide antibiotic, penam | antibiotic efflux | 99.56 | 100 |
| OCU473 | Strict | mtrA | | protein homolog model | resistance-nodulation-cell division (RND) antibiotic efflux pump | macrolide antibiotic, penam | antibiotic efflux | 99.56 | 100 |
| | Strict | Mycobacterium tuberculosis intrinsic murA conferring resistance to fosfomycin | C117D | protein variant model | murA transferase | fosfomycin | antibiotic target alteration | 97.32 | 98.56 |
| | Strict | RbpA | | protein homolog model | RbpA bacterial RNA polymerase-binding protein | rifamycin antibiotic | antibiotic target protection | 90.99 | 97.37 |
| OCU474 | Strict | mtrA | | protein homolog model | resistance-nodulation-cell division (RND) antibiotic efflux pump | macrolide antibiotic, penam | antibiotic efflux | 99.56 | 100 |
| | Strict | RbpA | | protein homolog model | RbpA bacterial RNA polymerase-binding protein | rifamycin antibiotic | antibiotic target protection | 90.99 | 97.37 |
| | Strict | Mycobacterium tuberculosis intrinsic murA conferring resistance to fosfomycin | C117D | protein variant model | murA transferase | fosfomycin | antibiotic target alteration | 97.32 | 98.56 |
| OCU475 | Strict | RbpA | | protein homolog model | RbpA bacterial RNA polymerase-binding protein | rifamycin antibiotic | antibiotic target protection | 90.99 | 97.37 |
| | Strict | Mycobacterium tuberculosis intrinsic murA conferring resistance to fosfomycin | C117D | protein variant model | murA transferase | fosfomycin | antibiotic target alteration | 97.32 | 98.56 |
| | Strict | mtrA | | protein homolog model | resistance-nodulation-cell division (RND) antibiotic efflux pump | macrolide antibiotic, penam | antibiotic efflux | 99.56 | 100 |
| OCU476 | Strict | RbpA | | protein homolog model | RbpA bacterial RNA polymerase-binding protein | rifamycin antibiotic | antibiotic target protection | 90.99 | 97.37 |
| | Strict | mtrA | | protein homolog model | resistance-nodulation-cell division (RND) antibiotic efflux pump | macrolide antibiotic, penam | antibiotic efflux | 99.56 | 100 |
| | Strict | Mycobacterium tuberculosis intrinsic murA conferring resistance to fosfomycin | C117D | protein variant model | murA transferase | fosfomycin | antibiotic target alteration | 97.32 | 98.56 |
| OCU477 | Strict | mtrA | | protein homolog model | resistance-nodulation-cell division (RND) antibiotic efflux pump | macrolide antibiotic, penam | antibiotic efflux | 99.56 | 100 |
| | Strict | RbpA | | protein homolog model | RbpA bacterial RNA polymerase-binding protein | rifamycin antibiotic | antibiotic target protection | 90.99 | 97.37 |
| | Strict | Mycobacterium tuberculosis intrinsic murA conferring resistance to fosfomycin | C117D | protein variant model | murA transferase | fosfomycin | antibiotic target alteration | 97.32 | 98.56 |
| OCU478 | Strict | RbpA | | protein homolog model | RbpA bacterial RNA polymerase-binding protein | rifamycin antibiotic | antibiotic target protection | 90.99 | 97.37 |
| | Strict | Mycobacterium tuberculosis intrinsic murA conferring resistance to fosfomycin | C117D | protein variant model | murA transferase | fosfomycin | antibiotic target alteration | 97.32 | 98.56 |
| | Strict | mtrA | | protein homolog model | resistance-nodulation-cell division (RND) antibiotic efflux pump | macrolide antibiotic, penam | antibiotic efflux | 99.56 | 100 |
| OCU479 | Strict | RbpA | | protein homolog model | RbpA bacterial RNA polymerase-binding protein | rifamycin antibiotic | antibiotic target protection | 90.99 | 97.37 |
| | Strict | Mycobacterium tuberculosis intrinsic murA conferring resistance to fosfomycin | C117D | protein variant model | murA transferase | fosfomycin | antibiotic target alteration | 97.32 | 98.56 |
| | Strict | mtrA | | protein homolog model | resistance-nodulation-cell division (RND) antibiotic efflux pump | macrolide antibiotic, penam | antibiotic efflux | 99.56 | 100 |
| OCU480 | Strict | mtrA | | protein homolog model | resistance-nodulation-cell division (RND) antibiotic efflux pump | macrolide antibiotic, penam | antibiotic efflux | 99.56 | 100 |
| | Strict | RbpA | | protein homolog model | RbpA bacterial RNA polymerase-binding protein | rifamycin antibiotic | antibiotic target protection | 90.99 | 97.37 |
| | Strict | Mycobacterium tuberculosis intrinsic murA conferring resistance to fosfomycin | C117D | protein variant model | murA transferase | fosfomycin | antibiotic target alteration | 97.32 | 98.56 |
| OCU481 | Strict | RbpA | | protein homolog model | RbpA bacterial RNA polymerase-binding protein | rifamycin antibiotic | antibiotic target protection | 90.99 | 97.37 |
| | Strict | mtrA | | protein homolog model | resistance-nodulation-cell division (RND) antibiotic efflux pump | macrolide antibiotic, penam | antibiotic efflux | 99.56 | 100 |
| | Strict | Mycobacterium tuberculosis intrinsic murA conferring resistance to fosfomycin | C117D | protein variant model | murA transferase | fosfomycin | antibiotic target alteration | 97.32 | 98.56 |
| OCU482 | Strict | RbpA | | protein homolog model | RbpA bacterial RNA polymerase-binding protein | rifamycin antibiotic | antibiotic target protection | 90.99 | 97.37 |
| | Strict | mtrA | | protein homolog model | resistance-nodulation-cell division (RND) antibiotic efflux pump | macrolide antibiotic, penam | antibiotic efflux | 99.56 | 100 |
| | Strict | Mycobacterium tuberculosis intrinsic murA conferring resistance to fosfomycin | C117D | protein variant model | murA transferase | fosfomycin | antibiotic target alteration | 97.32 | 98.56 |
| OCU483 | Strict | RbpA | | protein homolog model | RbpA bacterial RNA polymerase-binding protein | rifamycin antibiotic | antibiotic target protection | 90.99 | 97.37 |
| | Strict | mtrA | | protein homolog model | resistance-nodulation-cell division (RND) antibiotic efflux pump | macrolide antibiotic, penam | antibiotic efflux | 99.56 | 100 |
| | Strict | Mycobacterium tuberculosis intrinsic murA conferring resistance to fosfomycin | C117D | protein variant model | murA transferase | fosfomycin | antibiotic target alteration | 97.32 | 98.56 |
| OCU484 | Strict | mtrA | | protein homolog model | resistance-nodulation-cell division (RND) antibiotic efflux pump | macrolide antibiotic, penam | antibiotic efflux | 99.12 | 100 |
| | Strict | Mycobacterium tuberculosis intrinsic murA conferring resistance to fosfomycin | C117D | protein variant model | murA transferase | fosfomycin | antibiotic target alteration | 97.32 | 98.56 |
| | Strict | RbpA | | protein homolog model | RbpA bacterial RNA polymerase-binding protein | rifamycin antibiotic | antibiotic target protection | 90.99 | 97.37 |
| OCU485 | Strict | RbpA | | protein homolog model | RbpA bacterial RNA polymerase-binding protein | rifamycin antibiotic | antibiotic target protection | 90.99 | 97.37 |
| | Strict | Mycobacterium tuberculosis intrinsic murA conferring resistance to fosfomycin | C117D | protein variant model | murA transferase | fosfomycin | antibiotic target alteration | 97.32 | 98.56 |
| | Strict | mtrA | | protein homolog model | resistance-nodulation-cell division (RND) antibiotic efflux pump | macrolide antibiotic, penam | antibiotic efflux | 99.56 | 100 |
| OCU486 | Strict | mtrA | | protein homolog model | resistance-nodulation-cell division (RND) antibiotic efflux pump | macrolide antibiotic, penam | antibiotic efflux | 99.12 | 100 |
| | Strict | Mycobacterium tuberculosis intrinsic murA conferring resistance to fosfomycin | C117D | protein variant model | murA transferase | fosfomycin | antibiotic target alteration | 97.32 | 98.56 |
| | Strict | RbpA | | protein homolog model | RbpA bacterial RNA polymerase-binding protein | rifamycin antibiotic | antibiotic target protection | 90.99 | 97.37 |
| Toy194 | Strict | Mycobacterium tuberculosis intrinsic murA conferring resistance to fosfomycin | C117D | protein variant model | murA transferase | fosfomycin | antibiotic target alteration | 97.32 | 98.56 |
| | Strict | mtrA | | protein homolog model | resistance-nodulation-cell division (RND) antibiotic efflux pump | macrolide antibiotic, penam | antibiotic efflux | 99.56 | 100 |
| | Strict | RbpA | | protein homolog model | RbpA bacterial RNA polymerase-binding protein | rifamycin antibiotic | antibiotic target protection | 90.99 | 97.37 |
| Toy195 | Strict | RbpA | | protein homolog model | RbpA bacterial RNA polymerase-binding protein | rifamycin antibiotic | antibiotic target protection | 90.99 | 97.37 |
| | Strict | Mycobacterium tuberculosis intrinsic murA conferring resistance to fosfomycin | C117D | protein variant model | murA transferase | fosfomycin | antibiotic target alteration | 97.32 | 98.56 |
| | Strict | mtrA | | protein homolog model | resistance-nodulation-cell division (RND) antibiotic efflux pump | macrolide antibiotic, penam | antibiotic efflux | 99.56 | 100 |
