## Supplementary Tables for "Genomic features of *Mycobacterium avium* subsp. *hominissuis* isolated from pigs in Japan": Supplementary_Table 6_1st response.pdf

| Isolate | Contig | Locus tag | tRNA gene isotype synteny | Speies | Query cover | Identity | Accession |
| --- | --- | --- | --- | --- | --- | --- | --- |
| GM17 | Contig 1 | GBQ13_00450 - GBQ13_00590 | LLKKGCWPVMNYQQEFEINASHRRLITR | *Mycobacterium chimaera* strain MC045 genome assembly, plasmid: 2 | 64% | 85.97% | LT703506 |
| OCU479 | Contig 38 | GBP94_21805 - GBP94_21975 | TWLLKKGCPVMNYQQEFEIPASHRRLRI | *Mycobacterium chimaera* strain AH16 plasmid unnamed1 | 32% | 76.88% | CP012886 |
